# HCFC1 and YY1 mediate recruitment of COMPASS and Integrator to initiate X chromosome inactivation

**DOI:** 10.1101/2025.09.18.677040

**Authors:** Jeffrey Boeren, Beatrice F. Tan, Sarra Merzouk, Eveline Rentmeester, Rien van Haperen, Samuel J. Luchsinger-Morcelle, Cheryl Maduro, Richelle Rietdijk, Büşra Göynük, Wilfred F.J. van IJcken, Jeroen A.A. Demmers, Cristina Gontan, Hegias Mira Bontenbal, Joost Gribnau

**Author notes:** These authors contributed equally (Jeffrey Boeren, Beatrice F. Tan). These authors contributed equally (Cristina Gontan, Hegias Mira Bontenbal, Joost Gribnau).

## Abstract

The evolution of mammalian sex chromosomes has driven the emergence of mechanisms that balance X-linked gene dosage between male (XY) and female (XX) cells. In females, dosage compensation is achieved through X chromosome inactivation (XCI), initiated by upregulation of the long non-coding RNA *Xist*, which spreads in *cis*, recruiting chromatin modifiers to silence gene expression on one X chromosome. Here, we conducted a forward genetic screen and identified X-encoded Host Cell Factor 1 (HCFC1), a member of the COMPASS H3K4 methyltransferase complex, as a dose-dependent XCI-activator. HCFC1 loss results in genome-wide reduction of H3K4me3 at specific regulatory elements and downregulation of nearby genes, including *Xist*. We show that HCFC1 and YY1 are co-recruited genome-wide to gene regulatory elements. Mass spectrometry analysis confirmed an interaction of HCFC1 and YY1 and uncovered the Integrator complex as another prominent YY1 partner. YY1 depletion results in genome-wide loss of Integrator recruitment at gene regulatory elements and reduced expression of nearby genes, including *Xist cis*-regulatory genes *Jpx* and *Ftx*. These results highlight a co-regulatory role for HCFC1 in COMPASS recruitment and *Xist* activation, alongside YY1-mediated recruitment of Integrator to *Xist* regulatory elements and genes to activate female-exclusive XCI.

## Introduction

X chromosome inactivation (XCI) provides a paradigm for studying gene regulation in a chromosomal context. In mice and other mammals, random XCI is initiated in all female somatic cells shortly after implantation and can be modeled in female (XX) mouse embryonic stem cells (ESCs), which trigger XCI upon *in vitro* differentiation. This process is orchestrated by *Xist*, a long non-coding RNA (lncRNA) encoded within the X inactivation center (XIC), a genomic locus enriched for lncRNA genes that regulate *Xist* expression. Upon initiation of XCI, *Xist* coats the X chromosome in *cis* and recruits chromatin-modifying complexes to silence X-linked genes. Prior to and during early XCI, *Xist* transcription is repressed on the future active X by antisense transcription of the lncRNA *Tsix* (reviewed in ^1,2^).

Female-exclusive XCI is regulated by X-encoded activators of the XCI process directing dose-dependent activation of *Xist*^3,4^. The X-encoded E3 ubiquitin-ligase RNF12 was identified as an XCI-activator through dose-dependent degradation of pluripotency factor REX1^5,6^. YY1 and its homologue REX1 compete for binding at the same gene regulatory element of *Xist*, where REX1 recruitment prevents YY1-mediated activation of *Xist*^7^. Homozygous *Rnf12* knockout (KO) ESCs fail to initiate XCI during differentiation, whereas heterozygous loss of *Rnf12* leads to delayed XCI initiation with skewing towards inactivation of the X chromosome carrying the mutated *Rnf12* allele in *vitro*^8^. While the maternally inherited *Rnf12* allele is essential for imprinted XCI in extra-embryonic tissues in mice^9,10^, a conditional *Rnf12* KO mouse model showed that *Rnf12* is dispensable for the initiation of random XCI in the post-implantation epiblast^11^. Additionally, compound homozygous KO ESCs and mice of *Rnf12* and its primary target *Zfp42* (encoding REX1) display a partial rescue of the XCI phenotype, restoring it to near wild-type levels^10^.

These findings highlight the important role RNF12 plays in XCI regulation. Nonetheless, the fact that XCI is still initiated in *Rnf12*^+/−^ and *Rnf12*^−/−^:*Zfp42*^−/−^ differentiating female ESCs, but not in male ESCs, points to the presence of additional XCI activators. The lncRNAs *Jpx* and *Ftx*, respectively located 10 kb and 90 kb upstream of *Xist*, have been implicated as XCI-activators, playing important roles in *Xist* regulation (reviewed in^1,2^) However, deletion studies indicate that most of this activity is mediated in *cis*^8,12,13^. Also, the X-linked demethylase KDM5C has been described as an XCI-activator by catalyzing the conversion of H3K4me2/3 to H3K4me1 at the YY1-REX1 regulatory element of *Xist*^14^. Although *Kdm5c* KO female embryos are non-viable, *Xist* expression is unaffected in *Kdm5c* heterozygous KO females supporting the hypothesis that multiple XCI activators are at play^14^. Finally, the GATA transcription factor family has been implicated in XCI regulation; however, the only X-linked family member, *Gata1,* is not expressed in ESCs or the epiblast, indicating that other XCI-activators remain to be identified^15^.

Here, we used a forward genetic screen deleting large segments of the X chromosome to identify novel regulators of XCI and uncover the X-linked Host Cell Factor 1 (HCFC1) as a potent XCI activator. HCFC1, a component of the COMPASS H3K4 methyltransferase complex, promotes XCI by recruiting COMPASS to deposit H3K4me3 at the *Xist* regulatory element. Acute HCFC1 depletion during mouse ESC differentiation abolishes *Xist* and *Jpx* expression, preventing XCI. We found that genome-wide recruitment of HCFC1 is partly mediated by the transcription factor YY1, which we confirmed to physically interact with HCFC1. YY1 interactome analysis further identified the Integrator complex, a key mediator of transcriptional pause release, as a major YY1-associated partner. Loss of YY1 reduces Integrator binding at thousands of regulatory sites, including *Xist cis*-regulatory elements, without affecting H3K4me3 deposition. Together, these findings reveal a dual role for HCFC1 and YY1 in XCI initiation: HCFC1-mediated recruitment of COMPASS to establish H3K4me3 and YY1-mediated recruitment of Integrator to promote transcriptional activation of *Xist*.

## Results

### A forward genetic screen to identify novel XCI-activators

To identify new XCI-activators that act in parallel with *Rnf12* in the activation of XCI, we performed a CRISPR/Cas9-based screen deleting large regions of the X chromosome (Fig. 1a). We made use of female heterozygous *Rnf12^+/−^* F1 hybrid *Mus musculus castaneus* (Cast): *Mus musculus musculus* 129/Sv (129) ESCs providing a powerful SNP-based readout to differentiate the two X chromosomes. These cells also display complete skewed XCI of the *Rnf12* mutant X_129_ chromosome to retain one active copy of *Rnf12* X_cast_ throughout SC differentiation^8^. Due to this phenotype, large heterozygous deletions of the X_129_ chromosome are tolerated during differentiation except for deletions that affect XCI (Fig. 1a). The deletions generated covered all regions of the early evolutionary strata, which are expected to retain key regulatory elements and genes involved in XCI (Fig. 1b)^16–18^. Proper deletion of regions from the X_129_ chromosome was confirmed by PCR of allele-specific repeat amplifications, whole-genome sequencing (WGS) detecting SNPs between the X_129_ and X_Cast_ chromosomes, and DNA-FISH (Extended Data Fig. 1a-e and Extended Data Fig. 2a). Differentiation followed by *Xist* quantitative PCR (qPCR) analyses of at least two biological replicates per deletion indicated significant loss of *Xist* expression at day 8 of differentiation in deletion E (ΔE), located 29.5Mb centromeric to *Xist* (Fig. 1b,c). ΔE’s inability to upregulate *Xist* was subsequently confirmed by RNA fluorescent in situ hybridization (RNA-FISH) (Extended Data Fig. 2b-d). Deletion of regions harboring putative activators *Gata1* (located in ΔA and ΔAB) and *Kdm*5c (located in ΔK) did not result in a significant loss of *Xist* expression (Fig. 1c)^14,15^. To identify the gene(s) involved in XCI activation, we generated incremental and convergent deletions within ΔE (Fig. 1d) and determined the effect on *Xist* expression at day 4 and day 8 of ESC differentiation. ΔE2 and ΔE3 showed a reduction of *Xist* expression, unlike ΔE1, implying that our target gene(s) must be contained within ΔE2 and ΔE3 (Extended Data Fig. 2e). Next, we narrowed down the search by creating additional deletions within regions E2 and E3, confirmed by WGS and RNA-seq (Fig. 1e and Extended Data Fig. 1e) and discovered that ΔE2B and ΔE3B displayed a *Xist* phenotype at day 4 and day 8 of differentiation, but not ΔE2A and ΔE3A (Fig. 1f). RNA-seq analysis revealed that loss of an 80kb region encompassing *Hcfc1* and *Irak1* severely affects XCI at both analysed timepoints, whereas downregulation of pluripotency factors and upregulation of factors associated with ESC differentiation were broadly uncompromised compared to the parental cell line (Fig. 1e and Extended Data Fig. 2f). This analysis confirmed downregulation of *Xist* in ESC lines harboring ΔE2B and ΔE3B encompassing *Hcfc1* and *Irak1* (Fig. 1g,h) and as expected, impaired inactivation of X-linked genes (Extended Data Fig. 2g). Analysis of previously published datasets show that, similar to *Rnf12*, *Hcfc1* is upregulated between days 1 and 3 of ESC differentiation when XCI is initiated (Fig. 1i and Extended Data Fig. 2g) ^5,19^, whereas *Irak1* is not upregulated. HCFC1 has also been shown to interact with the crucial *Xist-*regulator YY1, pointing at *Hcfc1* as a candidate XCI activator ^7,20^.

**Figure 1:**
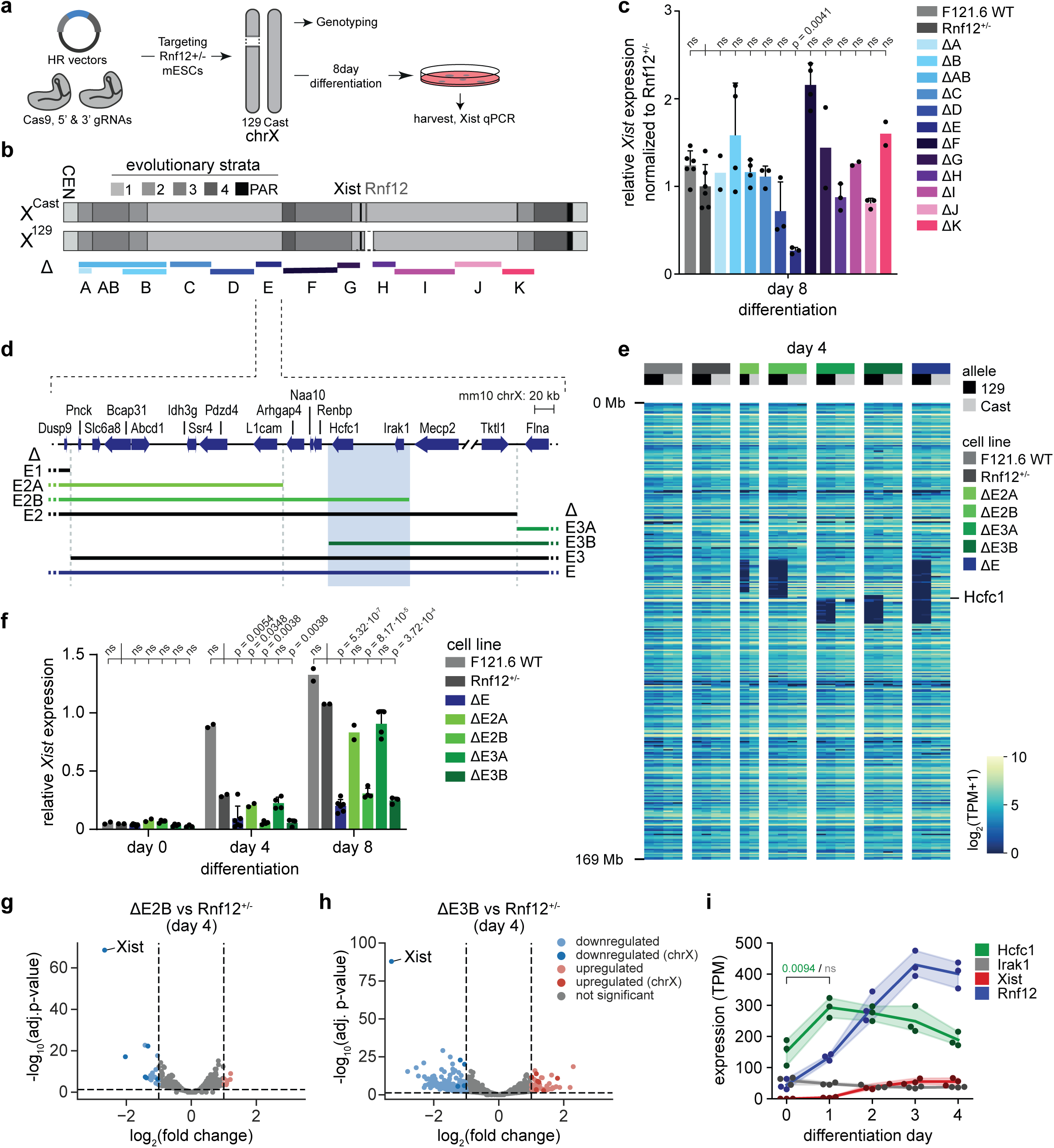
Large mega-domain deletions on the X chromosome identify HCFC1 as a potential XCI activator. **a**, Targeting and differentiation strategies. *Rnf12^+/−^* cells were transiently transfected with CRISRPR-Cas9 and gRNAs targeting different locations on the X chromosomes. Correctly targeted clones were differentiated for several days. **b**, X chromosome scheme depicting evolutionary strata 1, 2, 3, 4 and the pseudo-autosomal region (PAR)^17,18^. The locations of *Xist* and *Rnf12* are depicted with a black and grey line, respectively, while the deleted *Rnf12* allele on the 129 chromosome is also shown. The different deletions (ΔA to ΔK) are shown with different colors underneath. **c**, Relative *Xist* expression at day 8 of differentiation of WT, *Rnf12^+/−^* and the different *Rnf12^+/−^* cell lines with X chromosomal deletions, normalized to *Rnf12^+/−^*. Average ± SD, n=2 to 5 biological replicates. A one-sided Welch’s t-test was used to test significant differences between *Rnf12^+/−^* and WT or deletion cell lines. P-values were corrected for multiple testing with the Benjamini-Hochberg method. **d**, Zoom-in on a specific region of ΔE. The locations of different overlapping deletions (ΔE1, ΔE2, ΔE3) are shown with black lines; the smaller incremental deletions (ΔE2A, ΔE2B, ΔE3A, ΔE3B) are shown in green. The blue rectangle highlights the location of *Hcfc1* and *Irak1*. **e**, Expression heatmap of X-linked genes, sorted by genomic location and split by allele (129 or Cast) for 2 WT, 2 *Rnf12^+/−^*, 1 ΔE2A, 2 ΔE2B, 2 ΔE3A, 2 ΔE3B and 2 ΔE clones. Notice the deleted genes per clone on the 129 alleles. **f**, Relative *Xist* expression at day 0 and day 8 of WT, *Rnf12^+/−^*, ΔE and smaller ΔE deletion cell lines. Average expression ± SD; n=2 (WT, *Rnf12^+/−^* and ΔE2A lines), n=4 (ΔE2B, ΔE3A and ΔE3B lines) and n=5 (ΔE lines) biological replicates. A one-sided Welch’s t-test was used to test significant differences per day of differentiation between *Rnf12^+/−^* and WT or deletions cell lines. P-values were corrected for multiple testing with the Benjamini-Hochberg method. **g**, Volcano plot portraying gene expression changes between ΔE2B and *Rnf12^+/−^* cells. Downregulated and upregulated genes in ΔE2B are shown in blue and red, respectively. Notice *Xist* is depicted on the left, as the most significant downregulated gene. **h**, Same as **g** but for ΔE3B vs *Rnf12^+/−^* cells. **i**, Expression (TPM) of *Hcfc1* (green), *Irak1* (grey), *Rnf12* (blue) and *Xist* (red) in ΔXic_Cast_ ESCs from Pacini et al., 2021^19^. P-values were calculated between day 0 and 1 for *Hcfc1* (green) and *Irak1* (grey) using one-sided independent t-tests and corrected for multiple testing with the Benjamini-Hochberg method.

### *Hcfc1* is a dose-dependent activator of XCI

To test whether *Hcfc1* possesses XCI-activator activity, we restored *Hcfc1* expression in ΔE2B and ΔE3B ESC lines that showed an XCI phenotype by introducing a wild-type *Hcfc1* (129 origin) or a control transgene (Fig. 2a, Extended Data Fig. 3a). RT-qPCR analysis showed that *Xist* levels were rescued upon restoration of *Hcfc1* expression levels at day 4 of ESC differentiation in a transfected pool of cells, and in individual clones with a stably integrated transgene, confirming a dose-dependent regulatory role for *Hcfc1* in XCI (Fig. 2b,c, Extended Data Fig. 3b,c). To test whether HCFC1 is required for activation of XCI, we studied the effect of dTAG-13-mediated depletion of HCFC1 (Fig. 2d). Correct 3’ integration of a 2xFLAG-V5-tag combined with FKBP12^F36V^ in both *Hcfc1* (HCFC1-FKBP) alleles was confirmed by PCR (Extended Data Fig. 3d). HCFC1 is glycosylated and cleaved by OGT to form dimers that stay associated^21,22^. We confirmed loss of expression of both the N- and C-terminal domains after 48 hours of dTAG13 treatment (Fig. 2e, Extended Data Fig. 3e and 4e). Loss of HCFC1 occurs very rapidly, as examination of the C-terminal peptide showed a complete loss within 2 hours after addition of dTAG-13 (Fig. 2e). Depletion of HCFC1 during different time intervals of ESC differentiation resulted in a significant reduction of *Xist* expression (Fig. 2d,f). *Xist* RNA-FISH analysis confirmed loss of *Xist* expression and a severely reduced percentage of *Xist-*coated X chromosomes at day 3 of differentiation (Fig. 2g,h). Allele-specific RNA-seq analysis also showed that positive *Xist cis-*regulator *Jpx* was significantly downregulated, and *Ftx* showing a downregulation trend, whereas expression of pluripotency factors *Nanog*, *Pou5f1*, *Sox2* and *Rex1* remained unaffected, indicating that ESC differentiation proceeded normally in the absence of HCFC1 within the analysed time windows (Fig. 2i, Extended Data Fig. 3f). On the contrary, expression of several *trans*-regulators of *Xist*, including *Rnf12, Yy1*, *Kdm5c* and *Rif1* was significantly increased upon loss of HCFC1, suggesting a compensatory mechanism in *Xist* regulation (Extended Data Fig. 3g). dTAG-mediated loss of HCFC1 did not affect negative cis-regulators of *Xist*, such as *Tsx* and *Tsix* (Fig. 2i). Consistent with the observed substantial *Xist* downregulation, allele-specific expression analysis of X-linked genes indicated loss of XCI in HCFC1-depleted ESCs at day 3 of differentiation (Fig. 2j). To discern if this loss of silencing is due to *Xist* loss or HCFC1 loss, we removed HCFC1 and forced *Xist* expression in undifferentiated ESCs by means of a doxycyclin inducible promoter at the endogenous *Xist* locus. We observed that XCI still happens properly since X-linked *Rnf12* was silenced while a gene escaping XCI, *Kdm6a,* was not (Extended Data Fig. 3h-k). These findings identify HCFC1 as a dose-dependent activator of XCI and indicate that *Hcfc1* is required for upregulation of *Xist* and positive *Xist* regulatory genes.

**Figure 2:**
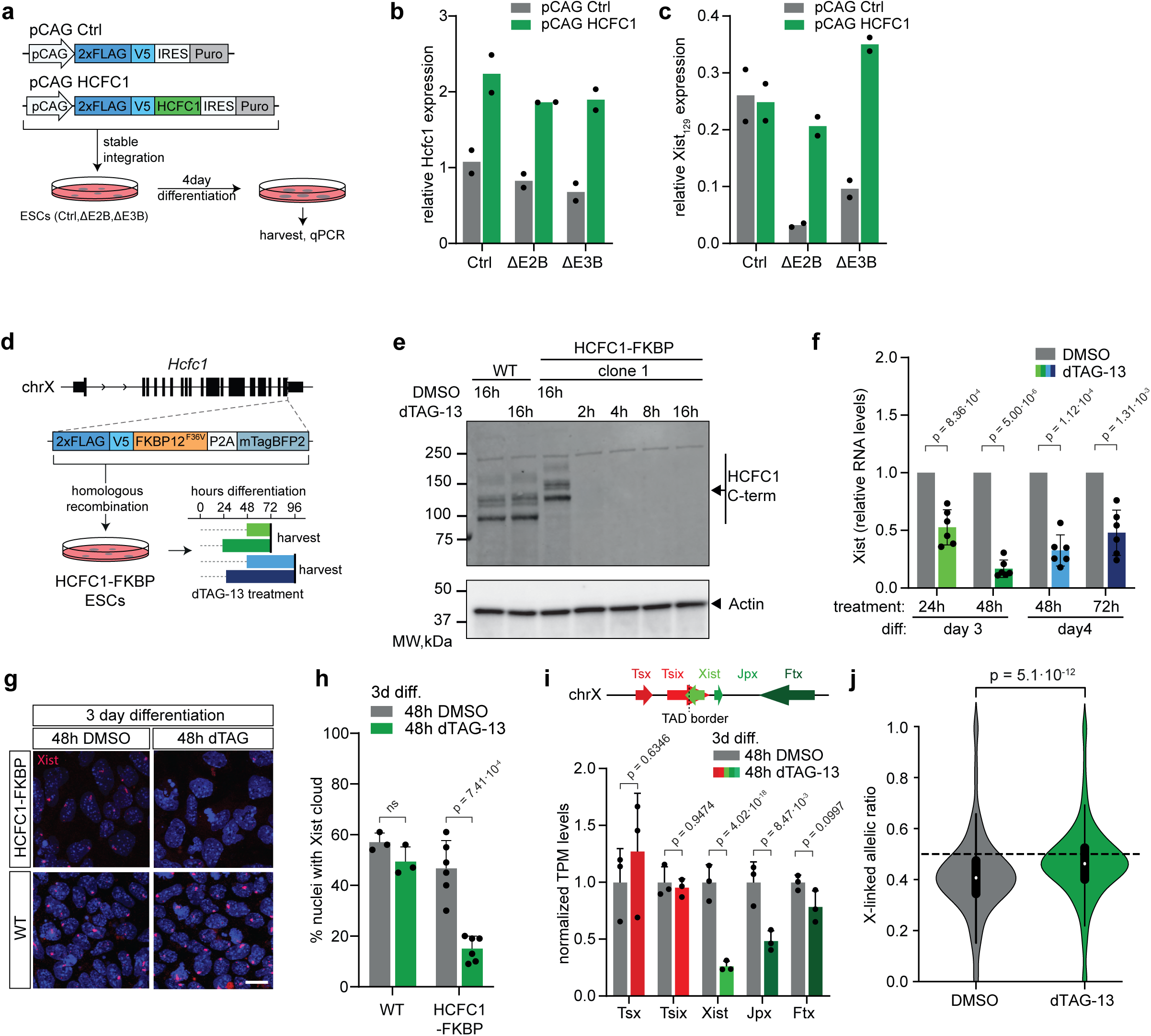
HCFC1 is an XCI activator. **a**, HCFC1 rescue and differentiation strategy of control (Ctrl; parental Rnf12^+/−^), ΔE2B and ΔE3B ESCs. **b**, RT-qPCR analysis of *Hcfc1* expression upon *Hcfc1* rescue in pools of transfected cells after 4 days of differentiation of in Ctrl (parental *Rnf12^+/−^*), ΔE2B and ΔE3B. The average of n=2 biological replicates is shown. **c**, RT-qPCR analysis of *Xist* expression in pools of transfected cells after 4 days of differentiation. The average of n=2 biological replicates is shown. **d**, 2xFLAG-V5-FKBP12^F36V^ tagging strategy of HCFC1 and differentiation schemes with different dTAG-13 treatment windows. **e**, Western blot of the C-terminal fragments of HCFC1 upon dTAG-13 treatment in undifferentiated ESCs. **f**, RT-qPCR analysis of *Xist* expression at different timepoints after start of dTAG-13 treatment and differentiation normalised to DMSO control; see Fig. 2a. A two-sided Welch’s t-test was used to assess significant differences between DMSO- and dTAG-13 treated cells. P-values were corrected for multiple testing using the Benjamini-Hochberg method. **g**, *Xist* RNA-FISH analysis in HCFC1-FKBP ESCs differentiated for 3 days and treated with dTAG-13 for the last 48h vs DMSO and parental ESCs. **h**, Quantification of **g**. Average ± SD, n=3-6 biological replicates, n=157-236. A two-sided Welch’s t-test was used to assess significant differences between DMSO- and dTAG-13 treated cells. P-values were corrected for multiple testing using the Benjamini-Hochberg method. **i**, Gene expression levels of several genes located in the XIC determined by RNA-seq analysis of HCFC1-FKBP ESCs differentiated for 3 days and treated with dTAG-13 for the last 48h. Shown in green are genes residing in the *Xist* TAD and involved in its activation. In red, genes located in the *Tsix* TAD and involved in *Xist* repression. Average TPMs normalized to 48h DMSO control ± SD, n= 3 biological replicates. P-values were calculated using DESeq2, which corrects for multiple testing using the Benjamini-Hochberg method. **j**, Violin plot of allelic ratios of X-linked gene expression in the DMSO and dTAG-13 conditions. The box plots display the median (black line), the interquartile range (box limits) and 1.5x of the interquartile range (whiskers). Dashed line indicates ratio of 0.5 as expected ratio without silencing. P-value was calculated using a two-sided Mann-Whitney U test.

### Overlapping genome-wide enrichment of HCFC1, YY1 and Integrator

HCFC1 is a member of the Set1A/B and MLL1/2 COMPASS complexes, mediating H3K4me3 deposition at gene regulatory elements which promotes transcription activation through Integrator complex recruitment and promoter-proximal eviction of paused RNA Pol II^23–26^. To mechanistically understand the role of HCFC1 in XCI and gene regulation in general, we re-analysed published ChIP-seq datasets examining the genome-wide distribution of HCFC1 and the COMPASS core member DPY30, as well as YY1, H3K4me3 and Ser-5 phosphorylated Pol II in undifferentiated mouse ESCs^25,27,28^. This analysis confirmed genome-wide co-enrichment of DPY30 with HCFC1 and their catalytic product H3K4me3 (Fig. 3a). Of 31,247 HCFC1 peaks in ESCs, 70.5% were found in genes, of which 48.7% were at promoters (Extended Data Fig. 4a). Within promoters, 81.5% of YY1 peaks overlapped with HCFC1 (Fig. 3b,c), confirming the previously described overlap in HCFC1 and YY1 binding to chromatin in human cells^20,29^. This analysis also shows that many HCFC1 peaks are not co-bound by YY1, indicating that other transcription factors might be recruiting HCFC1 to those sites.

**Figure 3:**
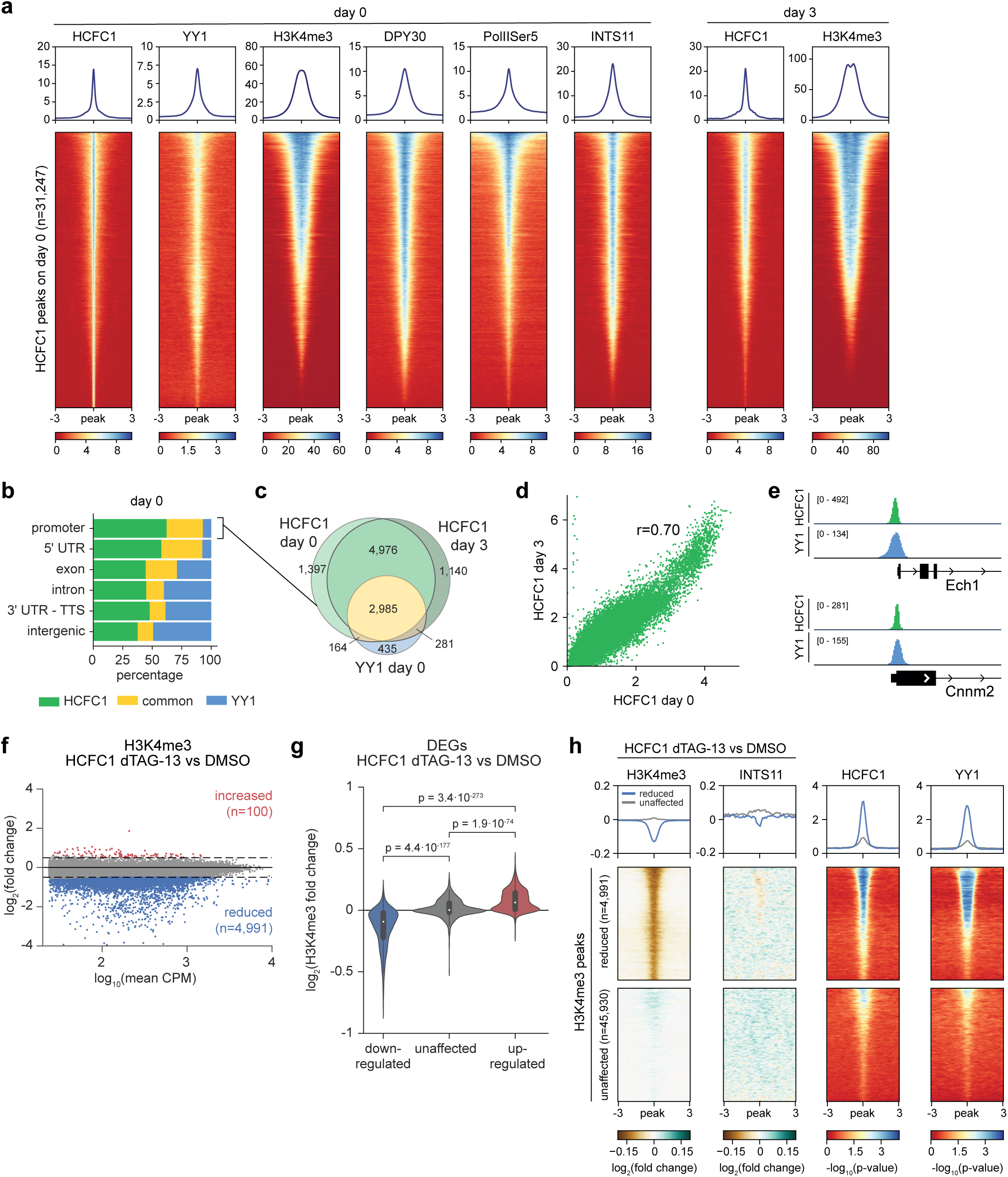
HCFC1 binding correlates with promoters, H3K4me3 deposition and YY1 binding. **a**, Heatmap of HCFC1, YY1, H3K4me3, DPY30, PolIISer5 and INTS11 enrichment at HCFC1 peaks in undifferentiated ESCs (left). The right panels show HCFC1 and H3K4me3 enrichment at day 3 of differentiation at HCFC1 peaks identified at day 0. **b**, HCFC1-only and YY1-only and shared HCFC1-YY1 peak distribution at different genomic locations at day 0. **c**, Venn diagram with the overlap of HCFC1 and YY1 peaks in promoter regions at day 0 and 3 of differentiation. **d**, Correlation between HCFC1 binding at day 0 and day 3 in promoter regions, with r representing the Pearson correlation coefficient. **e**, Genome browser views showing examples of HCFC1-YY1 overlapping peaks in promoter regions. **f**, MA plot depicting H3K4me3 enrichment changes upon dTAG-13-mediated removal of HCFC1. In red, peaks with increased H3K4me3 enrichment; in grey, unaffected peaks and in blue, peaks with reduced H3K4me3 enrichment. **g**, Violin plots showing H3K4me3 changes in promoter regions of differentially expressed genes from the RNA-seq comparison between HCFC1-FKBP dTAG-13 and DMSO. P-values were calculated using two-sided Mann-Whitney U tests, corrected for multiple testing using the Benjamini-Hochberg approach. **h**, Heatmaps showing differences in H3K4me3 and INTS11 enrichment at H3K4me3 peaks that are lost upon dTAG-13-mediated removal of HCFC1 and unaffected H3K4me3 peaks. For both H3K4me3 and INTS11, the fold change between dTAG-13 and DMSO treatment is shown. HCFC1 binding (day 3) and YY1 binding (day 0) are displayed on the right. For the unaffected peaks, a random subset of 4,991 peaks is shown.

To understand HCFC1-mediated regulation of *Xist*, we examined HCFC1’s distribution by ChIP-seq on day 3 of ESC differentiation when *Xist* expression is upregulated. For this, we generated 2xFLAG-V5-tagged HCFC1 (HCFC1-FLAGV5) female ESC lines through CRISPR/Cas9-mediated integration (Extended Data Fig. 4b). ChIP-seq analysis revealed 20,660 HCFC1 binding peaks, the majority of which (77.1%) were located within genes, of which 44.6% were at promoter regions (Extended Data Fig. 4a). Comparison of HCFC1 binding between day 0 and day 3 revealed only minor changes (Fig. 3c,d), and similarly, the overlap with YY1 binding sites was evident, with 84.5% of YY1 promoter peaks overlapping with HCFC1 peaks at day 3 of differentiation (Fig. 3c,e). As described for human cells, HCFC1 peaks were highly enriched at CpG islands and promoters (Extended Data Fig. 4c)(Michaud 2013). Moreover, HCFC1 peaks showed strong enrichment for the YY1 binding motif and motifs of other transcription factors implicated in HCFC1 recruitment, such as RONIN and ZNF143/STAF (Extended Data Fig. 4d)^29^. Similar to undifferentiated ESCs, H3K4me3 ChIP-seq analysis at day 3 of differentiation revealed a high level of overlap with HCFC1 peaks, consistent with HCFC1’s role as a member of the COMPASS complexes responsible for H3K4me3 catalysis (Fig. 3a).

Recent studies have indicated an important role for COMPASS-mediated H3K4me3 in recruitment of Integrator to target genes^25^. To investigate the role of HCFC1-COMPASS-mediated catalysis of H3K4me3 and subsequent recruitment of Integrator, we studied the presence and distribution of H3K4me3 and INTS11, a member of the endonuclease module of the Integrator complex, before and after dTAG-13-mediated depletion of HCFC1. Western blot analysis revealed a global decrease of H3K4me3 presence at day 3 of differentiation upon HCFC1 removal by dTAG-13 treatment (Extended Data Fig. 4e,f). ChIP-seq analysis identified a genome-wide reduction of H3K4me3 enrichment at 4,991 sites (9.8% of all 51,021 H3K4me3 peaks) upon HCFC1 removal, confirming an important role for HCFC1 in H3K4me3 deposition, although the loss was less prominent than reported for the depletion of DPY30 or RBPP5 (Fig. 3f, Extended Data Fig. 4g)^25^. As expected, correlation analyses between RNA-seq and ChIP-seq data showed that gene expression changes following HCFC1 depletion are positively associated with changes in H3K4me3 levels, (Fig. 3g, Extended Data Fig. 4h,i). Sites that displayed decreased H3K4me3 enrichment upon HCFC1 depletion did not exhibit or only exhibited a mild loss of INTS11 binding, indicating that INTS11 recruitment to those sites is predominantly H3K4me3 independent (Fig. 3h, Extended Data Fig. 4g,i). Interestingly, HCFC1 sites that showed reduced H3K4me3 deposition upon HCFC1 removal also displayed very strong YY1 binding (Fig. 3h). Our results indicate that HCFC1 plays a crucial role in H3K4me3 catalysis at HCFC1-YY1 co-bound sites, but that the presence of this histone modification is unrelated to the recruitment of Integrator.

### YY1-mediated recruitment of the COMPASS and Integrator complexes

Our results suggest a putative interaction between YY1 and HCFC1 in regulating H3K4me3 deposition. To test whether YY1 is indeed involved in recruitment of COMPASS to target sequences, we generated two female 2xFLAG-V5-tagged YY1 ES cell lines and performed FLAG pull-downs followed by mass spectrometry analysis (Extended Data Fig. 5a,b). Our data confirmed strong interactions with the INO80 chromatin remodeling complex, as previously described^30^ and with HCFC1 and COMPASS member RBBP5, and revealed weak interactions with other COMPASS complex subunits, likely reflecting indirect protein–protein interactions (Fig. 4a). Interestingly, we also identified most Integrator complex subunits (INTS1, INTS2, INTS4, INTS7, INTS8, INTS9, INTS11 and INTS12) as strong YY1 interactors, suggesting a role for YY1 in recruitment of Integrator to gene regulatory elements. This includes all subunits of the RNA endonuclease module of Integrator, responsible for cleavage of nascent RNA associated with paused RNA pol II^31^.

**Figure 4:**
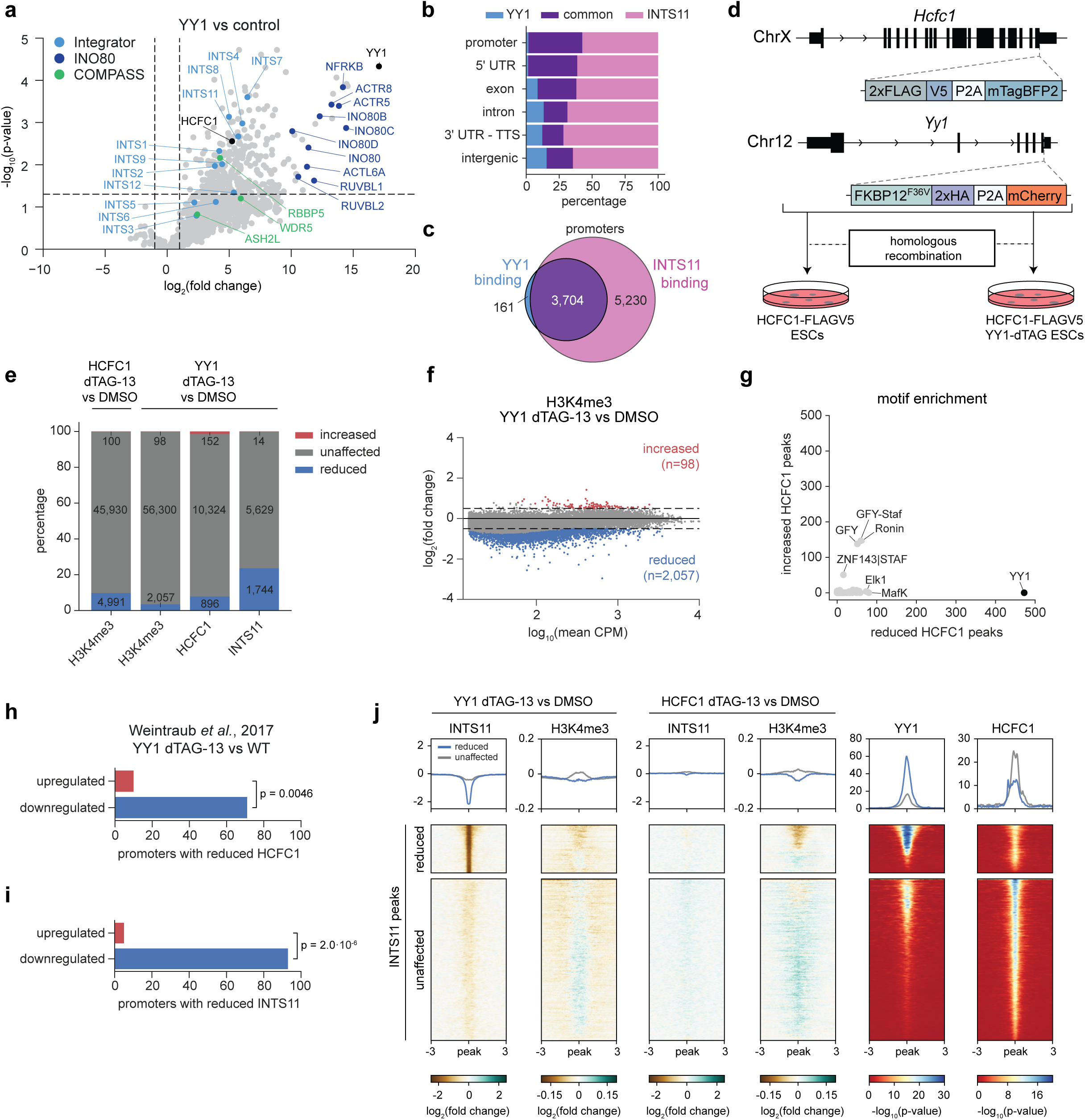
YY1 interacts with the Integrator complex. **a**, YY1 interactome identified by FLAG pull-down and mass spectrometry. Volcano plot showing proteins enriched in YY1 pull-downs compared to control. Known members of the Integrator (light blue), INO80 (dark blue), and COMPASS (green) complexes are highlighted. Selected proteins, including YY1 itself and HCFC1, are labeled in black. Dashed lines denote thresholds for significance (p < 0.05, log₂ fold change > 1). Data are based on two biological replicates for FLAG-YY1 and four biological replicates for the control. **b**, Percentages of shared YY1-INTS11, YY1-only or INTS11-only peaks at different genomic locations at day 0. **c**, Venn diagram showing the overlap of YY1 and INTS11 binding in promoter regions. **d**, Generation strategy of the FKBP12^F36V^-HA-Yy1:2xFLAG-V5-Hcfc1 compound knockin ESC lines by homologous recombination. **e**, Stacked barplot showing percentages of H3K4me3, HCFC1 and INTS11 peaks that significantly lost or gained enrichment or remained unchanged, upon HCFC1 or YY1 loss. **f,** MA plot depicting H3K4me3 enrichment changes upon dTAG-13-mediated removal of YY1. In red, peaks with increased H3K4me3 enrichment; in grey, unaffected peaks and in blue, peaks with reduced H3K4me3 enrichment. **g**, Scatter plot comparing motif enrichment between gained and lost HCFC1 peaks shown in **e**. Each point represents a motif, plotted by its −log10(p-value) in both analyses, and motifs with the lowest p-values in either condition are labeled. **h**, Barplot showing the number of up- and downregulated genes with reduced HCFC1 binding in the promoter region upon YY1 depletion. P-value was calculated using a Pearson’s chi-squared (χ²) goodness-of-fit test, based on the overall proportion of up- and downregulated genes. **i**, Same as **h**, but for promoter regions with reduced INTS11 binding. **j**, Heatmap showing changes in INTS11 and H3K4me3 enrichment upon dTAG-13-mediated depletion of YY1 or HCFC1, alongside YY1 and HCFC1 binding at INTS11 peaks. Peaks are grouped based on whether INTS11 enrichment is reduced (top) or unaffected (bottom) by YY1 depletion.

Interrogation of published ChIP-seq datasets of INTS11 distribution in undifferentiated ESCs indicates a high level of overlap between genome-wide distribution of YY1 and the Integrator complex (Fig. 4b, Extended Data Fig. 5c), with the immense majority (95.8%) of YY1 peaks at promoters overlapping with INTS11 (Fig. 4b,c). Indeed, overlapping YY1-INTS11 peaks are enriched for the YY1 motif while INTS11-only peaks are not (Extended Data Fig. 5d). These results suggest a dual role for YY1 in recruitment of both HCFC1 and Integrator complexes.

To functionally test whether HCFC1 and Integrator recruitment is YY1-dependent, we generated compound knock-in 2xFLAG-V5-Hcfc1:FKBP12^F36V^– 2xHA-Yy1 female (XX) ESC lines (HCFC1-FLAGV5/YY1-FKBP ESCs) and analysed the effect of dTAG-13-mediated depletion of YY1 on HCFC1 and Integrator recruitment to regulatory elements (Fig. 4d). Western blot analysis confirmed complete loss of YY1 within 2h of dTAG-13 treatment (Extended Data Fig. 5e). We then proceeded to differentiate these cells for 3 days and treated them with dTAG-13 for the last 2 days of differentiation (Extended Data Fig. 5e). HCFC1 ChIP-seq analysis at day 3 of differentiation revealed that HCFC1 enrichment at the great majority of its peaks does not change upon YY1 removal, indicating that HCFC1 recruitment is predominantly mediated by other transcription factor (TFs) (Fig. 4e). Nevertheless, 7.9% of HCFC1 peaks are YY1-dependent, and analysis of DNA binding motifs at HCFC1 peaks with decreased enrichment in the YY1 dTAG-13 condition compared to peaks with increased enrichment confirmed strong presence of the YY1 consensus binding motif (Fig. 4e,f). At 70 promoter-associated loci within this subset, loss of both YY1 and HCFC1 is associated with reduced gene expression (Fig 4h), indicating that YY1 recruits HCFC1 to a subset of promoters to activate their expression.

Since eight subunits of the Integrator complex were significantly pulled down by YY1^32^, we analysed INTS11 recruitment dependence on YY1. Loss of YY1 leads to reduction of INTS11 recruitment at 23.6% of all INTS11 binding sites genome-wide (Fig. 4e). INTS11 peaks with decreased enrichment in the YY1 dTAG-13 condition are strongly enriched for the YY1 binding motif (Extended Data Fig. 5f). Moreover, concomitant loss of YY1 and INTS11 at certain loci is associated with reduction in expression of 93 proximal genes such as *Cnnm2* and *Zfp37* at day 0^33^ and day 3 of differentiation (Fig. 4i, Extended Data Fig. 5g-k). Although western blot analysis showed a general decrease in H3K4me3 (Extended Data Fig. 5i,m), H3K4me3 ChIP-seq analysis indicated that 2,057 sites (3.5% of all 58,455 H3K4me3 peaks) showed a reduction of H3K4me3, a lower percentage compared to HCFC1 (Fig. 3f and Fig. 4e,i). Interestingly, most sites displaying a reduction of Integrator recruitment after YY1 depletion show persistent H3K4me3 enrichment (Fig. 4j). Importantly, at these sites INTS11 recruitment is HCFC1 independent (Fig. 4j), in line with our finding that dTAG-13-mediated depletion of HCFC1 revealed limited effects on INTS11 recruitment (Fig. 3h). These results highlight an important dual regulatory role for YY1 in genome-wide recruitment of COMPASS or Integrator to target genes to activate their expression.

### COMPASS- and Integrator-mediated regulation of XCI

To understand the XCI phenotype in *Hcfc1* mutant cells, we studied the role of HCFC1/COMPASS, YY1 and Integrator in XCI regulation. ChIP-seq analysis of HCFC1 and YY1 binding in day 0 ESCs showed clear HCFC1 and YY1 overlapping peaks around the *Xist, Jpx* and *Ftx* promoter and regulatory elements (Fig. 5a)^27,28^. Analysis of day 3 differentiated female ESCs revealed HCFC1 binding at the *Xist* regulatory element, as well as at regulatory elements of negative (*Tsix,*) and positive *(Jpx*, *Ftx)* regulators of *Xist* (Fig. 5a). Importantly, dTAG-13-mediated loss of HCFC1 or YY1 both leads to reduction of H3K4me3 at the *Xist* regulatory element accompanied by a reduction in *Xist* expression at day 3 of ESC differentiation (Fig. 2f and Fig. 5a,b). Integrator recruitment at the *Xist* regulatory element appears largely unaffected by HCFC1 and YY1 depletion and their concomitant loss of H3K4me3. *Jpx* expression is also significantly reduced upon HCFC1 or YY1 removal (Fig. 2i and 5b). In contrast to *Xist*, H3K4me3 deposition at the *Jpx* promoter remains unchanged in both conditions despite loss of HCFC1 recruitment at the *Jpx* promoter following YY1 depletion (Fig. 5a). Notably, INTS11 recruitment to the *Jpx* promoter is completely abrogated upon YY1 loss indicating a crucial role for YY1 in Integrator recruitment to the *Jpx* promoter. Although loss of HCFC1 results in a mild, non-significant reduction in *Ftx* expression, YY1 depletion causes a significant decrease in *Ftx* expression. In both cases, only INTS11 recruitment, and not H3K4me3 deposition, is affected (Fig. 2i and 5a,b), suggesting an indirect effect of HCFC1 on *Ftx* expression, possibly *via* loss of promoter co-regulation and/or mass action. While *Tsix* expression is affected by YY1 removal it is not affected by HCFC1 removal. Also, INTS11 recruitment to Tsix’ promoter is YY1-dependent and accompanied by a slight reduction in H3K4me3 deposition (Fig. 2i and 5a,b). Interestingly, the more severe effect of YY1 depletion on *Xist*, *Jpx* and *Ftx* compared to HCFC1 could be explained by the positive effect YY1 has on *Hcfc1* expression (Fig. 5b).

**Figure 5:**
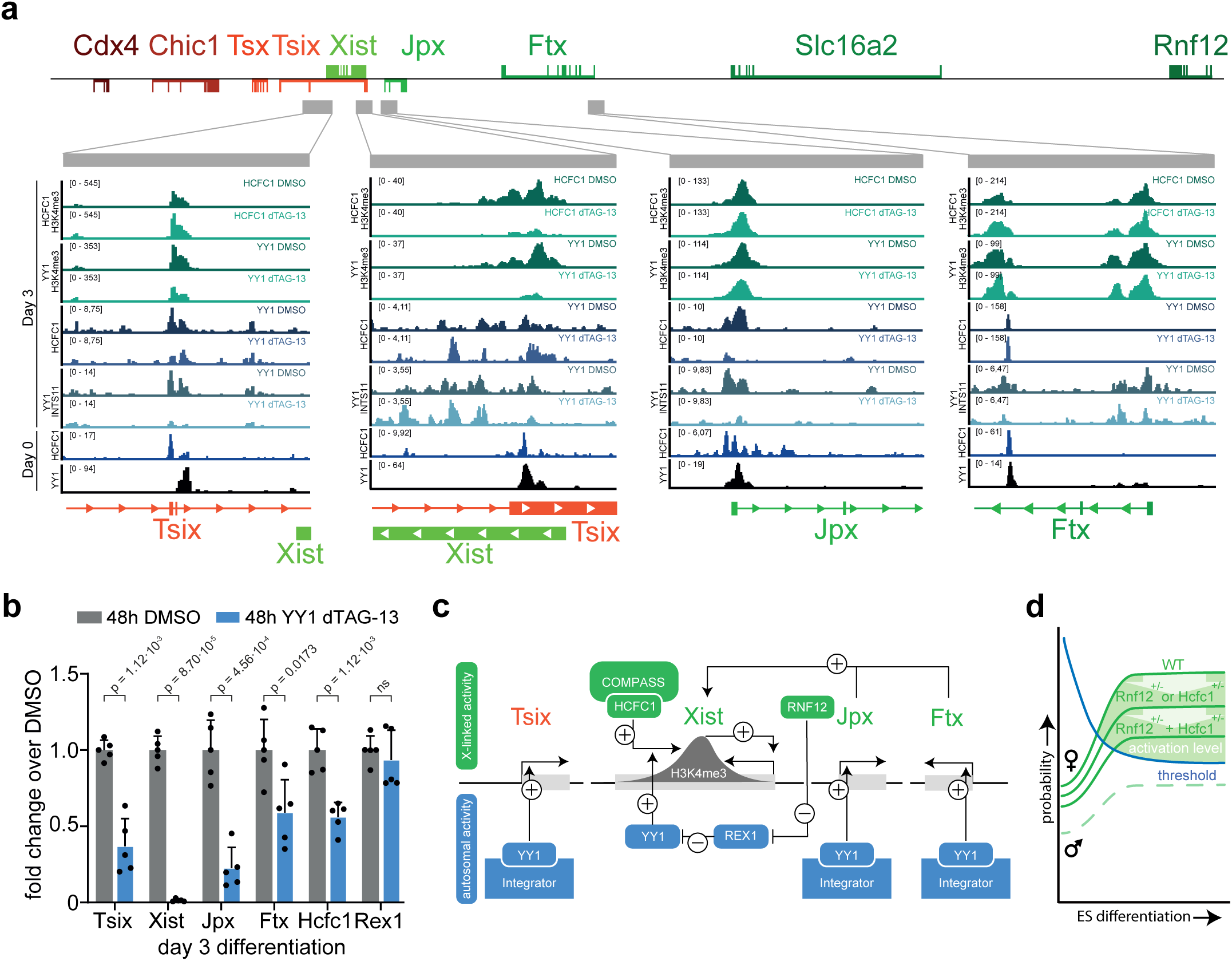
HCFC1 binds to and regulates the expression of several non-coding genes in the XIC. **a**, Genome browser views showing ChIP-seq tracks of H3K4me3 in HCFC1-depleted or YY1-depleted versus DMSO controls at day 3 of differentiation, HCFC1 ChIP-seq in YY1-depleted versus its DMSO control at day 3 of differentiation, INTS11 ChIP-seq in YY1-depleted versus its DMSO control at day 3, and YY1 and HCFC1 ChIP-seq in female ESCs at day 0 around the *Tsix*, *Xist*, *Jpx* and *Ftx* promoter regions. **b,** Relative expression of *Tsix, Xist*, *Jpx*, *Ftx, Hcfc1* and pluripotency factor *Rex1* at day 3 of differentiation of YY1-FKBP ESC cells normalised to DMSO control. A two-sided Welch’s t-test was used to test significant differences between DMSO- and dTAG-13 treated cells. P-values were corrected for multiple testing using the Benjamini-Hochberg method. **c,** Gene-regulatory network controlling *Xist* expression. RNF12 promotes YY1 binding by targeting the *Xist* repressor REX1 for proteasomal degradation. HCFC1/COMPASS and YY1 are both required for H3K4me3 catalysis at the *Xist* regulatory element. YY1 also recruits Integrator to activate transcription of *Xist* cis-regulatory genes within the XIC. **d**, Dose-dependent activation of *Xist* expression. Elevated levels of X-linked activators in XX cells raise the probability for *Xist* transcription initiation above threshold; this probability is reduced in *Rnf12*^+/−^ or *Hcfc*1^+/−^ cells and further diminished in double-heterozygous *Rnf12*^+/−^;*Hcfc*1^+/−^ cells.

These findings indicate distinct mechanisms in the regulation of XCI. HCFC1- and YY1-mediated H3K4 trimethylation is essential for *Xist* activation, with HCFC1 activity confined to *Xist*, while YY1 plays a dual role by recruiting the Integrator complex to both *Xist*-activating (*Jpx*, *Ftx*) and - repressing (*Tsix*) cis-regulatory elements. Interestingly, both HCFC1 and YY1 are required for H3K4me3 enrichment at the *Xist* locus, despite continued HCFC1 recruitment in the absence of YY1, suggesting an additional role for YY1 in catalyzing H3K4me3 at the *Xist* regulatory element.

## Discussion

Here, we provide evidence that HCFC1 plays a critical role in regulating XCI. Knockout and rescue experiments indicate that HCFC1 is a dosage-sensitive X-encoded activator of XCI involved in the regulation of *Xist* and *Xist* regulatory genes. Previous studies involving HCFC1 heterozygous knockout mice described an XCI phenotype with complete skewing of XCI towards inactivation of the mutant allele^34^. As described for *Rnf12*, this skewing phenotype could be explained by a continued requirement for *Hcfc1* during the XCI process but could also be explained by the essential function of HCFC1 in gene regulation in general and cell cycle control^34^. Unfortunately, early lethality of *Hcfc1*^−/−^ embryos due to imprinted XCI problems prohibited studying the role of HCFC1 in random XCI in the developing embryo.

Although the effect of the combined heterozygous deletions of large regions containing *Hcfc1* and *Rnf12* on *Xist* expression and XCI is severe, a low percentage of cells is still capable of initiating XCI upon ESC differentiation pointing to the presence of additional XCI-activators located in other regions and not detected in this study. Interestingly, our study did involve deletions encompassing genes previously implicated in regulating XCI including *Kdm5c* and *Gata1*, yet these deletions had no detectable effect on XCI in our system^14,15^. This result may be due to limited expression (*Gata1*) or sensitivity of our model system and readout (*Kdm5c*), which might not have been able to capture subtle effects on XCI.

HCFC1 is a member of the SET1A/B-COMPASS and MLL1/2-COMPASS complexes in mammals (reviewed in^35^). Our study confirms the importance of HCFC1-mediated recruitment of COMPASS to target sequences, H3K4me3 deposition, and proximal gene expression. However, in line with its non-core function within the complex, the effect of loss of HCFC1 on H3K4me3 deposition is less severe than depletion of core COMPASS subunits such as DPY30 or RBBP5^25,35^. Moreover, we show that Integrator recruitment to HCFC1-dependent H3K4me3 promoters is largely invariant, indicating other mechanisms at play to recruit Integrator to those sites. Interestingly, we noticed strong enrichment of most Integrator subunits in our YY1 pulldown studies, suggesting a direct role for YY1 in recruiting Integrator to target sites. Indeed, many YY1 sites showed reduced INTS11 recruitment upon YY1 removal and as with HCFC1, this seemed to be independent of H3K4me3, while the great majority of proximal promoters were downregulated. Also, although most HCFC1 binding seemed to be YY1 independent and thus mediated by other TFs, we observed a subset of YY1-dependent promoter-proximal HCFC1 sites that correlate with gene expression. Our study highlights the versatile role of YY1 in mediating recruitment of Integrator and HCFC1-COMPASS genome-wide. Although previous studies indicated a role for histone modification H3K4me3 in recruitment of the Integrator complex, our studies reveal a role for YY1 in recruitment of Integrator that seems independent of H3K4me3^25^.

By dissecting the contribution of YY1 and HCFC1 to XCI regulation, we show that YY1 and HCFC1 direct dose-dependent catalysis of H3K4 tri-methylation at the *Xist* regulatory element and activation of *Xist* transcription (Fig. 5c). Interestingly, although HCFC1 recruitment to this element appears unaffected in the absence of YY1, HCFC1 acute depletion still leads to loss of H3K4me3 suggesting a role for YY1 in recruiting COMPASS complexes independent of HCFC1, or indirectly by recruitment of other factors that facilitate H3K4me3 deposition. HCFC1 depletion also leads to expression reduction of the positive regulators of *Xist, Jpx*, *Ftx*, all located in the *Xist* topological associated domain (TAD). Since HCFC1 depletion does not lead to a loss of H3K4me3 at these co-activators, downregulation of *Jpx* and *Ftx* might be an indirect effect driven through loss of *Xist* co-activation. As HCFC1 has also been described as a member of the NSL complex, its loss may affect histone acetylation and thereby decrease gene expression of genes involved in XCI, a hypothesis that was not tested in the present study but warrants further investigation^36,37^. In contrast to the effects of HCFC1 loss, YY1 reduction decreases expression of genes in both the Xist-TAD and the neighbouring Tsix-TAD, which harbours negative regulators of *Xist*. Beyond its role in catalysing H3K4me3 at the *Xist* regulatory element, YY1 recruits both HCFC1 and INTS11 to the *Jpx* and *Tsix* promoters, and INTS11 alone to the *Ftx* regulatory element. Interestingly, YY1 depletion affects Integrator recruitment to these elements but loss of HCFC1 recruitment seems to have limited effect on H3K4me3 deposition pointing to an additional mechanism to recruit the core COMPASS complex independently of YY1 and HCFC1. Our results therefore indicate a role for YY1 in regulating *Xist* expression through H3K4me3 catalysis at its regulatory element, and regulating *Jpx, Ftx* and *Tsix* expression through recruitment of Integrator.

Our findings highlight the cooperative activities of HCFC1 and RNF12 in female exclusive initiation of XCI (Fig. 5d); through dose-dependent breakdown of REX1, RNF12 facilitates increased recruitment of YY1 to different regulatory elements at the XIC in female cells^7^. The effect on *Xist* expression is more pronounced in female cells likely through the dual action of YY1 on *Xist* and its positive regulators *Jpx* and *Ftx*, and *Hcfc1* itself. HCFC1 on the contrary, only appears to act through dose-dependent recruitment of COMPASS to the *Xist* regulatory region, where the combined action of YY1 and HCFC1 is required for catalysis of H3K4me3. In female cells, the increased activity of HCFC1, RNF12 and other XCI-activators amplifies the activation of *Xist* to sufficient levels to spread locally in *cis* and silence its negative regulator *Tsix.* Negative feedback will prevent inactivation of all X chromosomes except one. These findings highlight the complex and concerted action of RNF12 and HCFC1 in female exclusive initiation of XCI by dose-dependent action converging on the same XCI regulatory elements.

## Methods

### Cells and Cell Culture

#### Cell culture

F1 2-1 mESCs were grown on irradiated male mouse embryonic fibroblasts (MEFs) in medium containing DMEM (Gibco, 11995065) supplemented with 15% fetal calf serum, 100 U ml^−1^ penicillin, 100 µg mL^−1^ streptomycin (Sigma-Aldrich, P0781), 0.1 mM non-essential amino acids (Lonza, BE13-114E), 0.1 mM 2-mercaptoethanol (Gibco, 31350010) and 5.000 U mL^−1^ home-made LIF. Cells were passaged at least two times prior to differentiation. mESC were depleted of MEFs by pre-plating dissociated cells for 45 minutes at 37°C in non-gelatinized plates. The supernatant containing mESCs was counted and subsequently spun down for 5 min at 300 × g. Specific cell numbers were plated for different time points for monolayer differentiation in medium consisting of IMDM GlutaMAX™ (Gibco, 31980030) supplemented with 15% FCS, 0.1 mM NEAA, 100 U mL−1 penicillin, 100 µg mL−1 streptomycin, 23,75 µL L−1 monothioglycerol (Sigma-Aldrich, M6145) and 42 µg mL−1 ascorbic acid (Sigma-Aldrich, A92902). Degradation of HCFC1 or YY1 in mESCs or differentiation was achieved by refreshing the media with addition of 500 nM dTAG-13 (Tocris, 6605) or equivalent amounts of DMSO at the indicated time points.

#### Generating Rnf12 lines

Rnf12^+/−^ mESC lines were generated in F1 2–1 hybrid (129/Sv-Cast/Ei) mESC by CRISPR/Cas9 mediated targeting using different sets of gRNAs. The Rnf12^+/−^ ESC line Δ1 was generated by targeting mESCs with PX459 vectors with gRNAs targeting intron 2 and the 3’UTR using Gene Pulser 0.2 cm-gap cuvettes (Biorad, 1652086) at 118 kV, 1200 µF and ∞Ω in a Gene Pulser xCell electroporation system (Biorad). Twenty-four hours post-transfection, medium was refreshed and supplemented with 1 µg mL−1 puromycin (Sigma-Aldrich, P8833) for 36h. Drug-resistant clones were picked, expanded and characterized by PCR. Due to chromosomal instability upon inducing ΔJ, a second Rnf12^+/−^ line was generated (Rnf12^+/−^ Δ2). Rnf12^+/−^ Δ2 was generated by transfecting 2×1 µg PX459 alternative gRNA expressing vectors as well as 2 µg homology arm vector with lipofectamine 2000 (Invitrogen, 11668019). Twenty-four hours post-transfection, the medium was refreshed and supplemented with 1 µg mL−1 puromycin for 48h. Drug-resistant clones were picked, expanded and characterized by PCR.

#### Generating deletion lines

Deletion regions were designed so that genes within the same evolutionary strata and syntenic blocks were removed together^17,18^. As described previously, deletion lines were generated by transfecting 2×1 µg PX459 gRNA expressing vectors as well as 2 µg homology arm vector (containing a neomycin resistance cassette) with lipofectamine. 24h post-transfection, the medium was refreshed and supplemented with 1 µg mL−1 puromycin for 48h followed by 285 µg mL−1 Geneticin (G418) (Gibco, 11811-031) for 4-6 days for ΔA through ΔK. Combinations of 5′ and 3′ gRNAs were used to generate the incremental deletions ΔE1 to Δ E3 and ΔE2A to ΔE3B. No Geneticin selection was performed for the incremental deletions. Drug-resistant clones were picked, expanded, and characterized by microsatellite PCR. ΔA through ΔI and ΔK as well as all incremental deletions were generated in Rnf12^+/−^ Δ1. ΔJ was generated in Rnf12^+/−^ Δ2. No deletions were generated between ΔK and the telomere due to genomic instability.

#### Generating rescue lines

The coding sequence of *Hcfc1* was amplified from 129/Sv mESC cDNA and cloned into the pCAG-2×FLAG-V5-IRES-Puro expression vector. Rnf12^+/^, ΔE2B and ΔE3B mESCs were transfected with 2 µg of either the *Hcfc1* expression vector or an empty vector control, as described previously. 24h post-transfection, the culture medium was refreshed and supplemented with 1 µg mL⁻¹ puromycin for 5 days to select for stable integration. After two passages without selection, cells were split: one fraction was used to isolate individual clones, and the remaining pool was subjected to differentiation. Cells were harvested at day 4 of differentiation and analyzed by RT-qPCR.

#### Generation of knock-in ESC lines

Female mESCs harboring a heterozygous doxycycline responsive endogenous *Xist* promoter^38^ were used as a parental cell line for generation of endogenously tagged *Hcfc1* lines. In short, mESCs were transfected with gRNAs targeting the 3’ end and HCFC1-2×FLAG-V5-P2A-mTagBFP2 or HCFC1-2×FLAG-V5-FKBP^F36V^-P2A-mTagBFP2 constructs for homology-directed repair. 24 hours post-transfection cells were selected with medium supplemented with puromycin for 48 hours. Drug-resistant clones were picked on EVOS M5000 Imaging System (Thermo Scientific, AMF5000SV) for highest fluorescent intensity. To generate endogenous C-terminally tagged YY1 ESCs for dTAG-inducible degradation, HCFC1-2×FLAG-V5-P2A-mTagBFP2 cells were transfected with gRNAs targeting the 3’UTR and using a pAW62.YY1.FKBP.knock-in.mCherry vector^33^ (Addgene #10437) for homology-directed repair. Due to a SNP in the 3’UTR of *Yy1*, two separate gRNAs targeting either 129/Sv or Cast/Ei were selected. Drug-resistant clones were picked on EVOS M5000 Imaging System. All positive clones were expanded and screened for homozygous insertions flanking the integration site by PCR and western blotting.

Female ESCs expressing 2xFLAG-V5-YY1 were generated in a *Tsix-STOP* background by random integration of a pCAG 2xFLAG V5 YY1 construct. Cells were electroporated and subjected to puromycin selection 24 h later. After one week, individual clones were isolated and expanded. Transgene expression was confirmed by western blotting, and karyotypes were assessed. Two independent clones with normal karyotypes and YY1 expression comparable to endogenous levels were selected for FLAG pull down experiments.

### Western Blot

Nuclear protein extracts were performed as described previously^39^ with modifications. All buffers were supplemented with 1× protease inhibitors (Roche, 11836170001). Cells were harvested by scraping in cold PBS on ice and subsequently centrifuged for 5 min, 300 x g at 4°C. The pellet was resuspended in 5× pellet volume Buffer A (10 mM Hepes pH 7.6, 1.5 mM MgCl2 and 10 mM KCl, 0.5 mM DTT), incubated for 10 min at 4°C, vortexed for 30 sec and spun down at 900 x g, for 5 min, 4°C. The supernatant was transferred and the nuclear pellet was resuspended in 2× pellet volume Buffer C (20 mM Hepes pH 7.6, 25% glycerol, 420 mM NaCl, 1.5 mM MgCl2, and 0.2 mM EDTA, 0.5 mM DTT, 0.05% NP-40). After rotating the solution for 30 min, 4°C, it was subsequently spun down for 10 min at 18.000 x g, 4°C. The supernatant containing the nuclear fraction was collected and the concentration was determined using Nanodrop or The Pierce BCA Protein Assay Kit (Thermo Scientific, 23227). For western blot analysis, lysates were diluted 1:1 with 2× Laemmli Sample Buffer (0.125M Tris-HCl pH 6.8, 4% SDS, 20% (w/v) glycerol, 0.01% bromophenol blue, 150 mM DTT) and boiled for 5 min at 95°C. Equal concentration of protein lysates were loaded in the 4–15% Criterion™ TGX™ Precast Midi Protein Gel, 18-well (Biorad, #5671084) with Precision Plus Protein Dual Color Standards (Biorad, #1610374) and run in 1x Tris/Glycine/SDS buffer (Biorad, #1610772). Samples were transferred to a Trans-Blot Turbo PVDF membrane (Biorad, #1704157) using semi-dry transfer in the Trans-Blot Turbo Transfer System for 7 min at 2.5 A constant up to 25V. After transfer, membranes were blocked with 5% milk powder in 1×TBS (0.02M Tris-HCl, 0.15M NaCl, pH 7.6). The membrane was washed 3× with 1×TBST (0.02M Tris-HCl, 0.15M NaCl, 0.1% Tween-20, pH 7.6), incubated with the appropriate primary antibodies, HCFC1 Carboxy-terminal Antigen antibody (Cell signaling, 50708S, 1:1000), V5 antibody (Invitrogen, R96025, 1:5000), YY1 antibody (Santa Cruz, sc-7341, 1:200) and/or H3K4me3 antibody (Sigma-Aldrich, 05-1339, 1:1000). Membranes were washed 3× with 1× TBST and incubated with the appropriate secondary antibodies, IRDye 800CW Goat anti-Mouse IgG (LI-COR Biosciences, 926-32210), IRDye 800CW Donkey anti-Mouse IgG (LI-COR Biosciences, 926-32212) and/or IRDye® 680RD Donkey anti-Rabbit IgG (LI-COR Biosciences, 926-68073) and visualised on the Odyssey CLx Imager. Last, the membranes were incubated with β-Actin−Peroxidase (Sigma-Aldrich, A3854), stained with ECL Western blotting detection reagents (Cytyva, RPN2209) and visualised on the AI600 Imager.

### Expression analysis and RNA-seq

mESCs were depleted of MEFs as described above, supernatants were spun down and cell pellets were lysed. Differentiated cells were washed 1× in PBS and lysed directly in the culture dish prior to extraction. Total RNA was extracted using the ReliaPrep™ RNA Miniprep kit (Promega, Z6012) with increased DNase treatment (25 min at room temperature). Reverse transcription was performed on 500 ng RNA using Superscript III (Invitrogen, 18080093), random hexamers (Invitrogen, N8080127) and RNaseOUT (Invitrogen, 10777019). The cDNA was diluted 1:1 in water prior to amplification. RT-qPCRs, allele-specific RT-qPCRs and qPCRs were performed in triplicate in the GoTaq qPCR Master Mix (Promega, A6002) in a CFX384 Touch Real-Time PCR Detection System (BioRad, LJB22YE8Z). Allele-specific primer concentrations were determined and optimized using pure gDNA of 129/Sv and Cast/Ei mice. Rplp0 was used as reference gene for relative quantification of HCFC1-FKBP cells. Hist2h2aa1 was used as reference gene for all other qPCRs. All primers are included in Extended Data Table 1.

Sequencing libraries were prepared using the Illumina TruSeq stranded mRNA library preparation kit. Libraries from the deletion cell lines were sequenced on a HiSeq 2500 sequencer, generating single-end 50 bp reads. Libraries from the HCFC1 dTAG-13 and DMSO-treated samples were sequenced on an Illumina NextSeq 2000 sequencer, resulting in paired-end read clusters of 50 bases in length.

### RNA-seq analysis

RNA-seq reads were processed allele-specifically. First, we created an N-masked reference genome from mm10 using SNPsplit v0.3.4^40^, masking all SNPs between the 129/Sv and Cast/Ei strains based on the dbSNP v142 database^41^. Reads were trimmed using TrimGalore v0.6.7^42^ and aligned to the N-masked reference genome using HISAT2 v2.2.1^43^. Mapped reads were then assigned to either the 129/Sv, Cast/Ei, or unassigned BAM file based on the best alignment using SNPsplit (default settings with --paired for the paired-end data). All allele-specific and unassigned reads were merged into a composite BAM file using Samtools v1.10^44^.

Transcript-level counts were obtained from each BAM file using featureCounts v2.0.6^45^ with the Ensembl v98 gene annotations (-t exon -s 2). We normalized the raw counts to transcripts per million (TPM) using a custom Python script. Gene expression levels were visualized in bar plots, showing TPM values normalized to the average TPM of DMSO-treated or untreated control samples. To validate the deleted regions, log_2_(TPM+1) values were calculated per allele and visualized in a heatmap across all samples.

Differential expression analyses were performed using DESeq2 v1.30.1^46^ with parametric dispersion fitting. Fold changes were shrunk using the ashr v2.5-54 method^47^. Differentially expressed genes (DEGs) were defined as those with an adjusted p-value < 0.05 and an absolute log_2_(fold change) > 1. Results were visualized in volcano plots, with autosomal and X-linked DEGs coloured separately.

To assess X-chromosome silencing, we calculated allelic ratios for *Rnf12*^+/−^, ΔE, ΔE2B and ΔE3B as well as HCFC1 dTAG-13 and DMSO-treated samples. For each condition, genes with more than 20 allele-specific reads across all replicates were included. The allelic ratio was defined as:

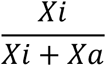

where Xi represents 129/Sv and Xa represents Cast/Ei. Allelic ratios of X-linked genes were visualized using violin plots combined with boxplots. Statistical significance was assessed using a two-sided Mann-Whitney U test (α < 0.05).

We reanalysed YY1-FKBP and wild-type RNA-seq data from GSE99521^33^ using the same allele-specific pipeline as described above, except that the N-masked reference genome was generated using SNPs between C57BL/6 and 129/Sv. Additionally, to assess expression dynamics of candidate genes during early differentiation, we analysed a bulk RNA-seq dataset from GSE151009^19^. Allele-specific count tables were downloaded and normalized to TPM values. For both ΔXic^B6^ and ΔXic^Cast^ cell lines, allelic expression of *Hcfc1*, *Irak1, Rnf12* and *Xist* was plotted over time in line plots, displaying the mean and 95% confidence interval. Statistical significance between day 0 and day 1 was assessed using one-sided independent t-tests, followed by multiple testing correction with the Benjamini-Hochberg method.

### RNA-FISH

Xist probes were made from a 19 Kb Xist/Tsix plasmid (p510) using the Nick Translation Kit (Abbott, 07J00-001) with red dUTPs (Abbott 02N34-050) according to the manufacturer’s instructions. Per coverslip, 3 µl of Xist probe was mixed with 1 µl salmon sperm DNA, 0.5 µl NaOAc 3M and 100 µl 100% EtOH and centrifuged 30 min, 12.800 rpm at 4°C. Pellet was washed with 500 µl cold 70% EtOH and let dry. 7 µl of formamide was added to the probe, incubated 10 min at 75°C and subsequently quenched on ice. 7 µl of hybridization buffer (20% Dextran sulfate, 2 mg/ml BSA, 20 mM VRC, 4×SSC), mixed and added to the coverslip.

ESCs were grown on gelatin coated coverslips at different time points for monolayer differentiation. Cells were fixed in 4% PFA in PBS for 10 min while rocking and rinsed 3× with PBS. Cells were subsequently permeabilized in 0.5% Triton, 20 mM VRC (NEB, S6639) in PBS for 4 min on ice. Cells were sequentially dehydrated by addition of 70, 90 and 100% ethanol, followed by airdrying. The probe, dissolved in hybridization buffer was added to the coverslip and incubated overnight at 37°C in humid chamber with 50% formamide in 2× SSC. After hybridization, coverslips were washed 3 × with 50% formamide in 2×SSC at 42°C, 2 × with 2×SSC at 42°C and once with 2 × SSC supplemented with 4 µM Hoechst (Thermo Scientific, 62249) before mounting with ProLong™ Gold Antifade (Invitrogen, P36930).

### WGS

ESCs were depleted of MEFs by pre-plating dissociated cells for 45 minutes at 37°C in non-gelatinized plates and subsequently grown for several days in a 6-well dish without MEFs. Cells were collected and lysed overnight in low-SDS lysis buffer (50 mM Tris-HCl pH 8.0, 100 mM EDTA, 0.2% SDS, 100 mM NaCl) with Proteinase K for Phenol-Chloroform DNA extraction. Following DNA extraction, 10 µg DNA was treated with 4 µg/ml RNase A treated for 1 hour at 37°C followed by clean up with the QIAamp DNA Micro Kit (QIAGEN, 56304). Sequencing libraries were prepared using the Nextera DNA Flex protocol from Illumina using 50 ng of genomic DNA and 5 cycles of PCR. Libraries were sequenced on an Illumina HiSeq 2500 sequencer, generating 50 bp single reads. For ΔE, over 50 million reads were obtained, while all other samples yielded at least 20 million reads.

### WGS analysis

Reads were trimmed using TrimGalore v0.6.7^42^ and aligned to the N-masked reference genome (see RNA-seq analysis) using Bowtie2 v2.5.1^48^. Mapped reads were assigned to either the 129/Sv or Cast/Ei BAM file using SNPsplit v0.3.4^40^, based on the best alignment. Allele-specific read counts were calculated for genomic windows of 100 kb using deepTools v3.5.5^49^ bamCoverage (-bs 100000 -of bedgraph --normalizeUsing None). For each window with more than 10 allele-specific reads, the log_2_(fold change) between the 129/Sv and Cast/Ei allele was computed. Statistical significance between both alleles was assessed using two-sided binomial tests with p=0.5, assuming equal read counts from both alleles in non-deleted regions. Resulting p-values were corrected for multiple testing using the Benjamini-Hochberg method. Data were visualized in scatter plots showing the log_2_(fold change) across the entire X-chromosome and within the 67-76.5 Mb region. For visualization, extreme log_2_(fold change) values were capped at ±2. Statistically significant windows (α < 0.05) were highlighted, as well as the window containing the *Hcfc1* locus.

### DNA FISH

BACs/Fosmids were selected inside (MeCP2; WI1-1364A8) and outside the deleted region (HPRT; RP23-412J16 and Xist; WI1-2363H9). Probes were labeled using the Nick Translation Kit (Abbott, 07J00-001) with red (Abbott 02N34-050) or green (Abbott 02N32-050) dUTPs according to the manufacturer’s instructions.

DNA FISH was performed as described previously^50^. Cells were blocked in metaphase by addition of 12 μl/ml KaryoMAX Colcemid (Gibco, 15210040) to the culture medium for 2 hours. Cells were trypsinized and resuspended in hypotonic buffer (0.075 M KCl) followed by fixation by addition of 3:1 methanol-acetic acid solution. Metaphase spreads were made by dropping fixed cells on coverslips and aged overnight at 37°C. Coverslips were equilibrated in 2×SSC (1× SSC: 0.15 M NaCl, 0.015 M sodium citrate), treated with RNAse A (1 µg/ml in 2×SSC) for 5 min at 37°C in humid chamber and washed 3× with 2×SSC at room temperature. Cells were permeabilized in pre-warmed 1% pepsin in 0.01M HCl solution for 7.5 min at 37°C. After permeabilization, cells were washed 2× with PBS and prefixed in 1% formaldehyde for 5 min at room temperature. Coverslips were washed 2× with PBS and sequentially dehydrated by addition of 70, 90 and 100% ethanol and airdrying.

5 µl of Nick-translated DNA probes was combined with 3 µg Mouse Cot-1 DNA (Invitrogen, 18440016) and 10 µg UltraPure™ Salmon Sperm DNA (Roche, 11467140001) and precipitated using NaAc and ethanol. The pellets were washed with 70% ethanol, spun down and air dried. The dried pellet was dissolved in formamide and denatured by incubating for 10 min at 75°C while shaking (1000 Rpm) and subsequently mixed 1:1 with cold 2×hybridization buffer (50% formamide, 2× SSC, 50 mM phosphate buffer (pH 7.0), 10% dextran sulfate and 1 ng/μl mouse Cot DNA). Following preparation of the probe mix, it was added to the coverslips and incubated on heatplate for 2 min at 75°C. Slides were transferred to a humid chamber with 50% formamide in 2× SSC and incubated overnight at 37°C. The next day, slides were washed 2× in 50% formamide in 2× SSC at 42°C, washed 2× in 2× SSC at 42°C, air dried and mounted with ProLong Gold Antifade with DAPI (Invitrogen P36931).

### ChIP-seq

ChIP-seq was performed as previously described^39^ with several modifications. Up to 1×10^8^ attached cells were fixed with 20mM disuccinimidyl glutarate (ThermoFisher, 20593-50MG) in PBS for 45min RT, washed twice with PBS followed by fixing in 1% PFA (Sigma-Aldrich, F1635-25ML) in PBS for 10min RT for the V5-HCFC1 and Integrator ChIPs, or only 1% PFA in medium for 10min RT for the H3K4me3 ChIPs (V5-HCFC1 ChIP was performed in FLAGV5-HCFC1 YY1-FKBP ESCs). After crosslinking, cells were quenched by addition of 125 mM glycine for 5 min. Cells were washed 1× with PBS before scraping in PBS with 1x protease inhibitors (Roche, 11836170001). From this point onwards, all buffers contained protease inhibitors (Roche, 4693132001) unless stated. Fixed cells were pelleted and subsequently resuspended in LB1 buffer (50 mM HEPES pH 8.0, 140 mM NaCL, 1 mM EDTA, 10% glycerol, 0.5% NP-40, 0.25% Triton X-100) and incubated for 10 min at 4 °C while rotating, then in LB2 buffer (10 mM Tris-HCl pH 8.0, 200 mM NaCl, 1 mM EDTA, 0.5 mM EGTA) and last in LB3 buffer (10 mM Tris-HCl pH 8.0, 100 mM NaCl, 1 mM EDTA, 0.5 mM EGTA, 0.1% Sodium Deoxycholate, 0.5% N-laurolylsarcosine). DNA was sheared using a Bioruptor Pico (Diagenode, B01060001) for 10 cycles of 30 sec on and 30 sec off (for single crosslinked samples) or 14 cycles of 30 sec on and 30 sec off (for double crosslinked samples) to an average fragment length of 100-300 bp. 1% triton X-100 was added to the sheared chromatin and incubated for 10 minutes at 4°C and spun down for 10 min at 4500 x g. Chromatin was aliquoted, flash frozen in liquid nitrogen and stored at −80°C until further use. Different amounts of chromatin were used for the H3K4me3 ChIPs (25 ug), HCFC1-V5 and INTS11 ChIPs (150 ug). Each chromatin sample was diluted to 500 ul using ChIP dilution buffer (1.1% Triton X-100, 0.01% SDS, 167 mM NaCl, 16.7 mM Tris-HCl and 1.2 mM EDTA) and 1% of it was substracted for INPUT and frozen it at −20°C. A specific amount of antibody was added (Ab; 2 ug H3K4me3 Ab, Cell Signaling Technologies, 9751S [C42D8]; 10 ug INTS11 Ab, Bethyl Laboratories, A301-274A; 7ul V5 Ab, Invitrogen R96025) to each chromatin sample and incubated with rotation overnight at 4°C. The next day, 30 ul Dynabeads protein G beads were washed (Invitrogen, 10004D) 2x with 500 ul ChIP dilution buffer. Chromatin with antibody was added to the beads and incubated 2h at 4°C. Afterwards, by magnetic separation, beads were washed 5 min 4°C 3x with Low salt wash buffer (20 mM Tris-HCl, 2 mM EDTA, 1% Triton X-100, 150 mM NaCl, 0.1% SDS), 1x with High salt wash buffer (20 mM Tris-HCl, 2 mM EDTA, 1% Triton X-100, 500 mM NaCl, 0.1% SDS) and 1x with LiCl wash buffer (10 mM Tris-HCl, 1 mM EDTA, 0.25 M LiCl, 1% NP-40, 1% deoxycholate). Beads were finally washed with TE without protease inhibitors. 200 ul of ChIP elution buffer (0.1 M NaHCO_3_, 10 mM EDTA and 1% SDS) were added to the beads with 22 ml proteinase K (10 mg/mL) and 5 ml RNase (10 mg/mL), and shaken at 1,000 rpm on an orbital thermal-mixer for 2 h at 37°C and finally at 65°C overnight. DNA was then cleaned using a Zymo ChIP DNA clean concentrator kit (Zymo Research, D5202) and eluted in 12 ul. 0.5 ul was used to confirm ChIP efficiency. The concentration of the ChIP sample was measured, and sequencing libraries were prepared using a ThruPLEX DNA-seq Kit (Takara Bio, R400675) following manufacturer’s instructions. Libraries were sequenced on an Illumina NextSeq 2000 sequencer, generating 50 bp paired-end reads.

### ChIP-seq analysis

Reads were trimmed using TrimGalore v0.6.7^42^ and aligned to the N-masked reference genome (see RNA-seq analysis) using Bowtie2 v2.5.1^48^. For datasets containing *Drosophila melanogaster* spike-ins, reads were also aligned to the dm6 genome, and those with higher alignment scores to dm6 than to mm10 were separated into a dedicated FASTQ file. All remaining reads were assigned to either the 129/Sv, Cast/Ei, or unassigned BAM file based on the best alignment using SNPsplit (--paired). Allele-specific and unassigned reads were merged into a composite BAM file using Samtools v1.10^44^.

Peak calling was performed on the composite BAM files using MACS3 v3.0.1^51^ with the callpeak function (-f BAMPE), comparing ChIP samples to their corresponding input controls. Bigwigs with signal tracks were generated using MACS3 bdgcmp with the ppois method. Peaks were annotated using the annotatePeaks.pl script from Homer v4.11^52^, and motif enrichment analysis was conducted with findMotifsGenome.pl (-size 200). Small annotation categories were merged for clarity: 3’ UTR and TTS were combined into “3’ UTR - TTS”, and exon and non-coding into “exon”.

Several ChIP-seq datasets were reanalysed using the same approach as described above. Reads from a YY1 ChIP-seq dataset in day 0 ESCs were obtained from GSE240684^27^ and analysed allele-specifically. In addition, we downloaded ChIP-seq data for H3K4me3, DPY30, PolIISer5, and INTS11 in day 0 ESCs from GSE181714^25^, as well as a HCFC1 ChIP-seq dataset in day 0 ESCs from GSE49847^28^. As these datasets were generated from non-hybrid cells, reads were mapped to the mm10 genome, and the resulting BAM files were used as composite BAM files for all subsequent analyses.

### Overlap between peak locations

To assess overlap between HCFC1 binding and other chromatin features, HCFC1 day 0 peaks were used as a reference to evaluate enrichment of HCFC1 (day 3), H3K4me3 (day 0 and 3), YY1, DPY30, PolIISer5, and INTS11 (all day 0). Heatmaps were generated using deepTools v3.5.5 with computeMatrix (reference-point mode, averageTypeBins mean, 10bp bins, ±3kb) followed by plotHeatmap.

Overlap between HCFC1 (day 0 and 3) and YY1 (day 0) peaks was quantified using intervene v0.6.5^53^ venn (--type genomic --save-overlaps). Resulting BED files for each overlap category were annotated with Homer annotatePeaks.pl. The proportion of peaks per annotation category was calculated for each overlap group and visualized in stacked bar plots. To specifically evaluate promoter-associated binding, peaks were filtered for those located in promoter regions, and their overlap was visualized in a Venn diagram using matplotlib-venn v1.1.1. Overlap between YY1 and INTS11 binding on day 0 and 3 was assessed following the same approach.

Correlation between HCFC1 binding at day 0 and day 3 was further assessed using deepTools multiBigwigSummary in BED-file mode, with a BED file containing all TSSs ±500 bp in mm10. Counts (obtained with outRawCounts setting) were compared in a scatter plot using log1p-transformed values. Pearson correlation was calculated using the plotCorrelation function with the removeOutliers option.

### Differential peak calling in H3K4me3 enrichment

To detect global changes in H3K4me3 levels, *Drosophila melanogaster* spike-ins were incorporated into all H3K4me3 samples. The number of reads mapping better to dm6 than mm10 (see ChIP-seq analysis) were counted and used to scale CPM-normalized tracks. Initial visualization of these tracks revealed large variability between replicates (data not shown). Consequently, the spike-in normalization was excluded from the final analysis. Instead, all H3K4me3 tracks were normalized using the ChIPseqSpikeInFree v1.2.4 method^54^. Composite BAM files were processed to compute read coverage in sliding windows, which were normalized to CPM and used to calculate scaling factors for each sample.

Differential peak calling between dTAG-13 and DMSO conditions was performed using DESeq2 v1.30.1^46^. Peaks identified by MACS3 in each condition were merged using BEDTools v2.31.1 merge^55^, and the resulting BED file was converted to SAF format. Read counts per merged peak were quantified with featureCounts v2.0.6^45^ and provided to DESeq2. The estimated size factors were adjusted using the scaling factors computed by ChIPseqSpikeInFree. A Wald test was performed to compare dTAG-13 and DMSO conditions, and fold changes were shrunk using the ashr v2.5-54 method^47^. Differential peaks were defined as those with an adjusted p-value < 0.05 and an absolute log_2_(fold change) > 1.

Tracks for each sample were scaled using the size factors computed by ChIPseqSpikeInFree. Average tracks per condition were generated with deepTools bigwigAverage. Moreover, tracks with log_2_(fold change) between dTAG-13 and DMSO were computed using deepTools bigwigCompare. To show HCFC1- and YY1-associated changes in H3K4me3 enrichment, we generated heatmaps with deepTools computeMatrix (reference-point mode, averageTypeBins mean, 100bp bins, ±3kb) and plotHeatmap. Differential peaks with significantly reduced H3K4me3 signal (as identified by DESeq2) were compared to unaffected peaks, which were randomly downsampled to match the number of reduced peaks. Both log₂(fold change) tracks and DMSO tracks were plotted for comparison.

### Differential peak calling in HCFC1 and INTS11 binding

Changes in HCFC1 and INTS11 binding in YY1-FKBP cells after dTAG-13 treatment were evaluated using MACS3 bdgdiff. The genome-wide number of unique and overlapping peaks were visualized in Venn diagrams using matplotlib-venn. Peaks from each category were annotated using the annotatePeaks.pl script from Homer v4.11 (Heinz et al. 2010), after which small annotation categories were merged for clarity: 3’ UTR and TTS were combined into “3’ UTR - TTS”, and exon and non-coding into “exon”. Motif enrichment analyses were conducted on each peak category separately with Homer findMotifsGenome.pl (-size 200). Enrichment results for known motifs were compared using scatter plots, where p-values for each motif were plotted between two categories, and motifs were coloured by TF family.

### Correlation between RNA-seq and ChIP-seq in HCFC1-FKBP cells

To correlate RNA-seq changes between HCFC1-FKBP dTAG-13 and DMSO treatments with changes to H3K4me3 and INTS11 ChIP-seq signal under the same conditions, we first selected DEGs from the RNA-seq analysis (adjusted p-value < 0.05, absolute log_2_(fold change) > 1). To allow comparison with unaffected expressed genes, we matched each gene in the downregulated group to a gene in the non-significant group with a similar expression level. This was done using the NearestNeighbors algorithm (n_neighbors=1) from scikit-learn v1.5.1 (Pedregosa et al., 2011), applying the fit and kneighbors functions. To avoid duplicate matches, each selected nearest neighbor was removed from the pool after selection. This resulted in a non-significant gene subset of equal size and similar expression distribution as the downregulated group. Finally, we visualized fold changes in H3K4me3 and INTS11 enrichment following dTAG-13 treatment, as well as baseline enrichment of HCFC1, YY1, H3K4me3, and INTS11 at TSSs of upregulated, downregulated, and matched unaffected genes.

We quantified H3K4me3 changes in the promoter region using deepTools multiBigwigSummary in BED-file mode, with a bed file containing regions spanning TSS + 1kb, based on the enrichment pattern observed in the ChIP-seq heatmap. The resulting mean signal change per promoter region was visualized in violin plots for each gene category separately. Statistical significance between groups was assessed using a two-sided Mann-Whitney U test, with p-values corrected for multiple testing using the Benjamini-Hochberg method (α < 0.05).

### Correlation between YY1-FKBP dTAG-13 vs DMSO RNA-seq and INTS11 ChIP-seq

To assess the relationship between transcriptional changes upon YY1 degradation and alterations in INTS11 binding, we compared RNA-seq data from YY1-FKBP dTAG-13 and DMSO-treated samples with changes in the INTS11 ChIP-seq signal under the same conditions. INTS11 peaks that were significantly reduced upon YY1 dTAG-13 treatment (see ‘Changes in HCFC1 and INTS11 binding’) were intersected with YY1 ChIP-seq peaks using intervene venn (type genomic, save-overlaps). Overlapping peaks were annotated with Homer annotatePeaks.pl, and genes with shared peaks located within 1 kb of a transcription start site (TSS) were selected. The resulting gene list was compared to the set of DEGs from the RNA-seq analysis. The number of up- and downregulated genes among these shared targets was visualized in a bar plot. Statistical significance was assessed using a Pearson’s chi-squared (χ²) goodness-of-fit test, comparing the observed number of up- and downregulated genes to the expected distribution based on the overall proportions of up- and downregulated genes.

### Genome browser views

Genome browser views were created using IGV v2.18.2^56^. ChIP-seq signal tracks were generated using the bdgcmp function in MACS3 (see ChIP-seq analysis). Log_2_(fold change) tracks comparing dTAG-13 and DMSO conditions were computed based on CPM-normalized coverage for HCFC1 and INTS11 binding, or ChIPseqSpikeInFree-scaled CPM-normalized tracks for H3K4me3 (see “Differential peak calling in H3K4me3 enrichment”) using deepTools bigwigCompare.

### Mass Spectometry

#### Nuclear extracts preparation

All subsequent steps for nuclear extract preparation and FLAG-purification were performed at 4°C using buffers supplemented with 1× protease inhibitors (Roche, 4693132001) and 15 µM MG-132 proteasome inhibitor (Sigma, C2211). Pre-chilled buffers and equipment were used throughout, and low protein-binding Falcon and Eppendorf tubes were employed to minimize sample loss. One day prior to harvest, two independently generated FLAG-V5-YY1 and 4 control mESC lines were trypsinized, MEFs were removed by pre-plating for 45 min in non-gelatinized culture dishes. The resulting mESCs were cultured without MEFs for 16h before processing. Cells were washed with cold PBS, collected by scraping on ice in PBS+ (1×PBS, 0.5 mM DTT), transferred to a Falcon tube and pelleted by centrifugation at 1500 rpm for 5 min. Pellets were resuspended in 2 pellet volumes of PBS+ and centrifuged at 1500 rpm for 5 min. The pellets were resuspended in 5 pellet volumes of buffer A (10 mM Hepes pH 7.6, 1.5 mM MgCl₂, 10 mM KCl, 0.5 mM DTT), incubated on ice for 10 min and centrifuged at 3000 rpm for 5 min. The supernatant was removed, and the pellets were resuspended in 2 pellet volumes of buffer A. Cells were lysed with 10 strokes of a Dounce homogenizer (type A pestle) on ice. The homogenate was transferred to Falcon tubes and centrifuged at 3000 rpm for 5 min. The supernatant, containing the cytoplasmic fraction, was discarded. The nuclear pellet was resuspended in 1.5 pellet volumes of Buffer C (20 mM Hepes pH 7.6, 20% glycerol, 420 mM NaCl, 1.5 mM MgCl₂, 0.2 mM EDTA, 0.5 mM DTT) and homogenized with 10 strokes of a Dounce homogenizer (type B pestle) on ice. The nuclear suspension was transferred to a Falcon tube, incubated while rotating for 30 min and centrifuged at 14000 rpm for 15min. The supernatant, containing the nuclear fraction, was collected and protein concentration was determined using a Nanodrop.

#### FLAG pull down

Protein concentrations of nuclear extracts were adjusted to the lowest concentration using buffer C. For each sample, 1 mL nuclear extract was diluted with 1 mL buffer C-0 (20 mM Hepes pH 7.6, 20% glycerol, 1.5 mM MgCl₂, 0.2 mM EDTA), to reduce the salt concentration to approximately 150 mM NaCl. Samples were incubated on ice for 15 min, centrifuged at 4000 rpm for 15 min, and the supernatant was transferred to a new tube. Benzonase nuclease (200 U; Novagen, 70664) was added to prevent RNA- or DNA-mediated interactions. Anti-Flag M2 Affinity Gel (Sigma-Aldrich, A2220) beads slurry (120 µl per sample) was washed four times with 1 ml Buffer C-150 (20 mM Hepes pH 7.6, 20% glycerol, 150 mM KCl, 1.5 mM MgCl₂, 0.2 mM EDTA, 0.02% NP40, 0.5 mM DTT) by centrifugation at 1000 rpm for 1 min. Nuclear extracts were added to the washed beads, resuspended, and incubated with gentle rotation for 3 h. Anti-FLAG-beads were pelleted by centrifugation at 2000 rpm for 2 min and washed five times with 1 mL buffer C-100 (20 mM Hepes pH 7.6, 10% glycerol, 100 mM KCl, 1.5 mM MgCl2, 0.2 mM EDTA, 0.02% NP40, 0.5 mM DTT). Each wash was performed with 5 min rotation (10 min for the final wash), followed by centrifugation at 2000 rpm for 2 min. The bound fraction was eluted by resuspending the Anti-FLAG-beads with 70 µL 0.2 mg/mL FLAG tripeptide peptide (Sigma, F4799) in buffer C-150 (without NP40), incubating for 15 min on ice, and centrifuging at 2000 rpm for 2 min. The supernatant was collected as the eluted fraction, and the elution step was repeated four times. An aliquot of the eluted fraction was analyzed by Western blot to confirm the presence of FLAG-V5-YY1 protein. Eluted fractions were combined and centrifuged at 2000 rpm for 2 min to remove residual Anti-FLAG beads. Protein eluates were concentrated by precipitation with trichloroacetic acid (w/v) / 0.1% sodium deoxycholate (w/v). Pellets were resuspended in 2× Laemmli sample buffer, boiled for 5 min at 95°C, separated by SDS–PAGE on 10% NuPAGE Bis-Tris Gel (Invitrogen, NP0315BOX) and stained using a colloidal blue staining kit (Invitrogen, LC6025).

#### Sample preparation

Complete lanes of SDS–PAGE gels were excised and divided into smaller slices. Proteins were reduced with 10 mM dithiothreitol (DTT) for 1 h at room temperature, alkylated with 55 mM chloroacetamide (CAM) for 1 h in the dark, and digested overnight at room temperature with sequencing-grade trypsin (1:100, w/w; Roche). Peptides were desalted using Sep-Pak tC18 Vac cartridges (Waters) and eluted with 80% acetonitrile (AcN).

#### MS analysis

Peptides were analyzed by nanoflow LC–MS/MS using an EASY-nLC system (Thermo) coupled to a Fusion Tribrid Orbitrap mass spectrometer (Thermo) operating in positive ion mode and equipped with a nanospray source. Peptide mixtures were first trapped on a ReproSil C18 reversed-phase column (Dr Maisch GmbH; 1.5 cm × 100 μm, packed in-house) at a flow rate of 8 μL/min, then separated on a ReproSil C18 reversed-phase column (Dr Maisch GmbH; 15 cm × 50 μm, packed in-house) using a linear gradient of 0–80% buffer B (0.1% FA, 80% v/v AcN) over 70 or 120 min at a constant flow rate of ∼200 nL/min. The column effluent was directly sprayed into the ESI source of the mass spectrometer. Spectra were acquired in profile mode with an MS1 resolution of 70,000 (AGC target: 3E6) across an m/z range of 350–1700. Data-dependent acquisition (Top15) was performed with HCD fragmentation, using an isolation window of 3.0 m/z and a normalized collision energy of 26–28%. MS2 spectra were recorded in the ion trap. Singly charged precursors were excluded, dynamic exclusion was set to 20 s, and the intensity threshold was set to 8.0E3.

#### Data analysis

Raw data were processed with the MaxQuant software suite (version 1.6.10.43)^57^ with label-free quantification (‘LFQ’) and intensity-based absolute quantification (‘iBAQ’) enabled. A false discovery rate (FDR) of 1% was applied at both protein and peptide levels, with a minimum peptide length of seven amino acids. MS/MS spectra were searched with the Andromeda search engine against the Mus musculus UniProt database (October 2019 release), concatenated with reversed sequences and a contaminant database. Up to two missed cleavages were allowed. Carbamidomethylation of cysteine was set as a fixed modification, and methionine oxidation as a variable modification. A minimum of two peptides, including at least one unique or razor peptide, was required for positive identification. Only unique and razor peptides that were non-modified, methionine oxidized, or N-terminally acetylated were used for quantification.

#### Statistical analysis

Quantitative proteomics data were analyzed using Perseus (https://www.maxquant.org/perseus/;^57^). The proteingroups.txt output file was filtered to remove contaminants, reverse hits, and proteins identified by a single tryptic peptide. Bona fide interactors were identified by two-sample, two-tailed t-tests with permutation-based FDR correction. iBAQ values were log₂-transformed, and missing values were imputed from a normal distribution (width = 0.3, shift = 1.8). Putative interactors (t-test difference > 1) were retained for further experimental validation.

## Data availability

All raw and processed high-throughput sequencing data (RNA-seq, WGS and ChIP-seq) generated in this study have been submitted to the NCBI Gene Expression Omnibus (GEO) under accession number (https://www.ncbi.nlm.nih.gov/geo/query/acc.cgi?acc=GSE306492, token: kjczegkqdfcnzql) and <u>GSE282784</u>. Moreover, RNA-seq datasets from <u>GSE99521</u> and <u>GSE151009</u>, and ChIP-seq datasets from <u>GSE240684</u>, <u>GSE181714</u> and <u>GSE49847</u> were reanalysed. The mass spectrometry proteomics data have been deposited to the ProteomeXchange Consortium *via* the PRIDE partner repository with the dataset identifier PXD060869 (https://www.ebi.ac.uk/pride/login, token: WIW4YME97KAe).

## Acknowledgements

We thank Mariana Pelicano de Almeida and Debbie van den Berg for the INTS11 ChIP protocol and feedback. We also thank Stanley van Herk for access to the Bioruptor Pico. Finally, we thank Danilo Remmers and Koert van den Bergh for technical assistance.

## Author contribution

J.B., B.T., S.M., C.G., H.M.-B. and J.G. conceptualised and designed experiments. J.B, S.M., E.R., R.v.H., S.J.L.-M., C.M., R.R., B.G., C.G. and H.M.-B. performed experiments and analysed data. B.T. performed all bioinformatic analyses. J.D. and W.v.I performed the mass spectrometry and sequencing, respectively. J.B., B.T., C.G., H.M.-B. and J.G wrote the manuscript and all authors edited it. J.B., S.M., C.M. and B.G. generated all cell lines. C.G. performed the pulldowns. J.B., S.M. and R.v.H. genotyped cell lines. J.B., S.J.L.-M. and H.M.-B. performed ChIP-seq experiments. J.B., E.R. and R.v.H. performed WB and RNA-FISH experiments. C.G., H.M.-B. and J.G. jointly supervised the project.

## Additional information

### Supplementary information

The online version contains supplementary material available at XXX.

### Materials and correspondence

All requests for materials and related questions should be addressed to Hegias Mira-Bontenbal and Joost Gribnau.

---

**Extended Data Figure 1:**
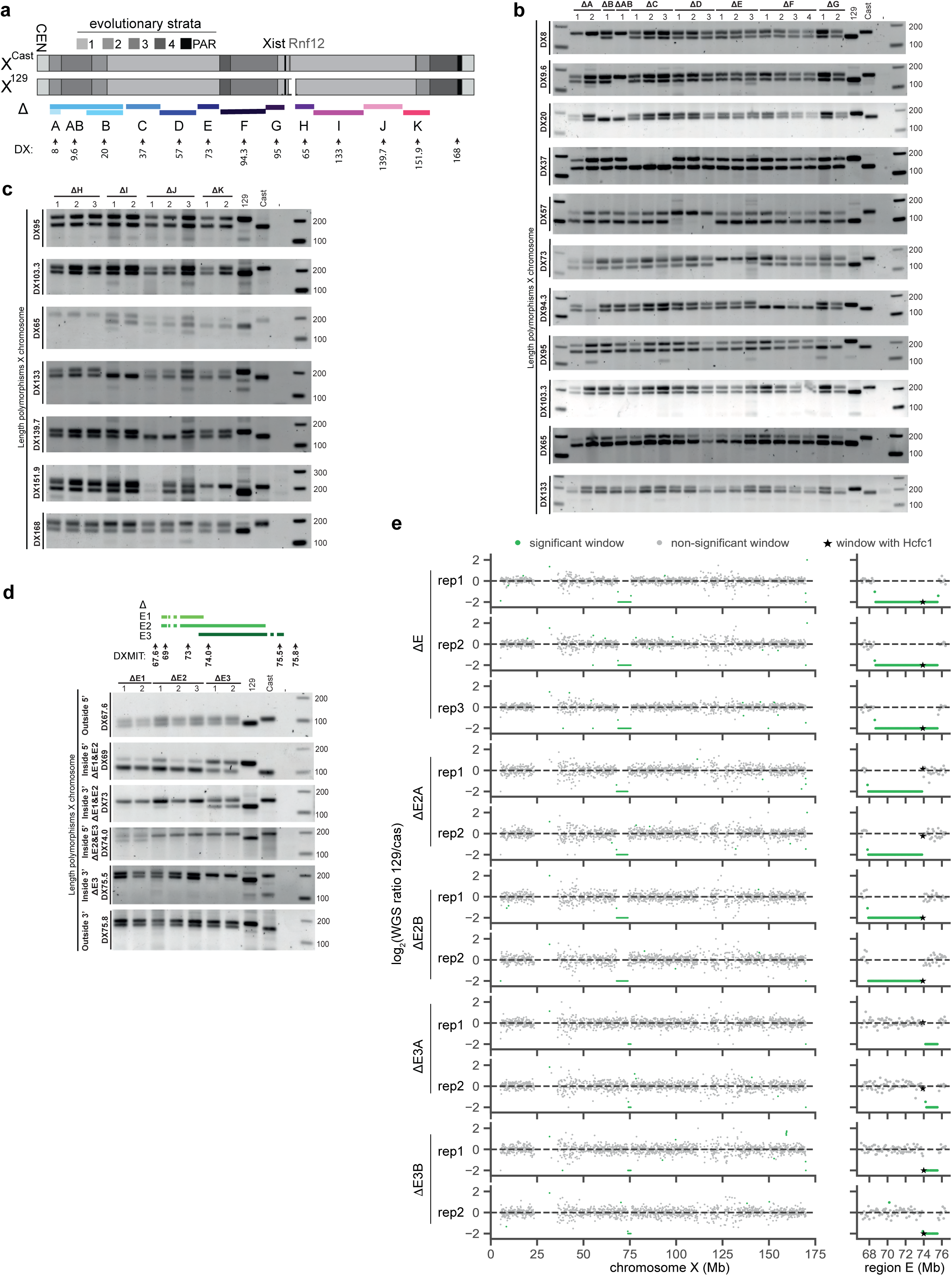
Megadomain deletion confirmation by PCR and WGS. **a**, Overview of the locations of the different deletions generated in this study. Different X-linked length polymorphisms between 129 and Cast are shown (DX amplicons). **b-c,** PCR analysis of different DX amplicons for the different deletions. **d**, Location of the different DX amplicons with respect to the deleted regions, and length polymorphism analysis inside and around several incremental deletions within region E. **e**, Scatter plots of allelic ratios along the X-chromosome from shallow whole-genome sequencing for different ΔE, ΔE2A, ΔE2B, ΔE3A and ΔE3B clones (left). Y-axis represents the log_2_(fold change) between the 129 and Cast alleles for each window. Windows with significant signal differences between alleles are highlighted in green. The zoomed-in plots on the right show the region around 67-77 Mb, with the *Hcfc1*-containing window indicated by a star.

**Extended Data Figure 2:**
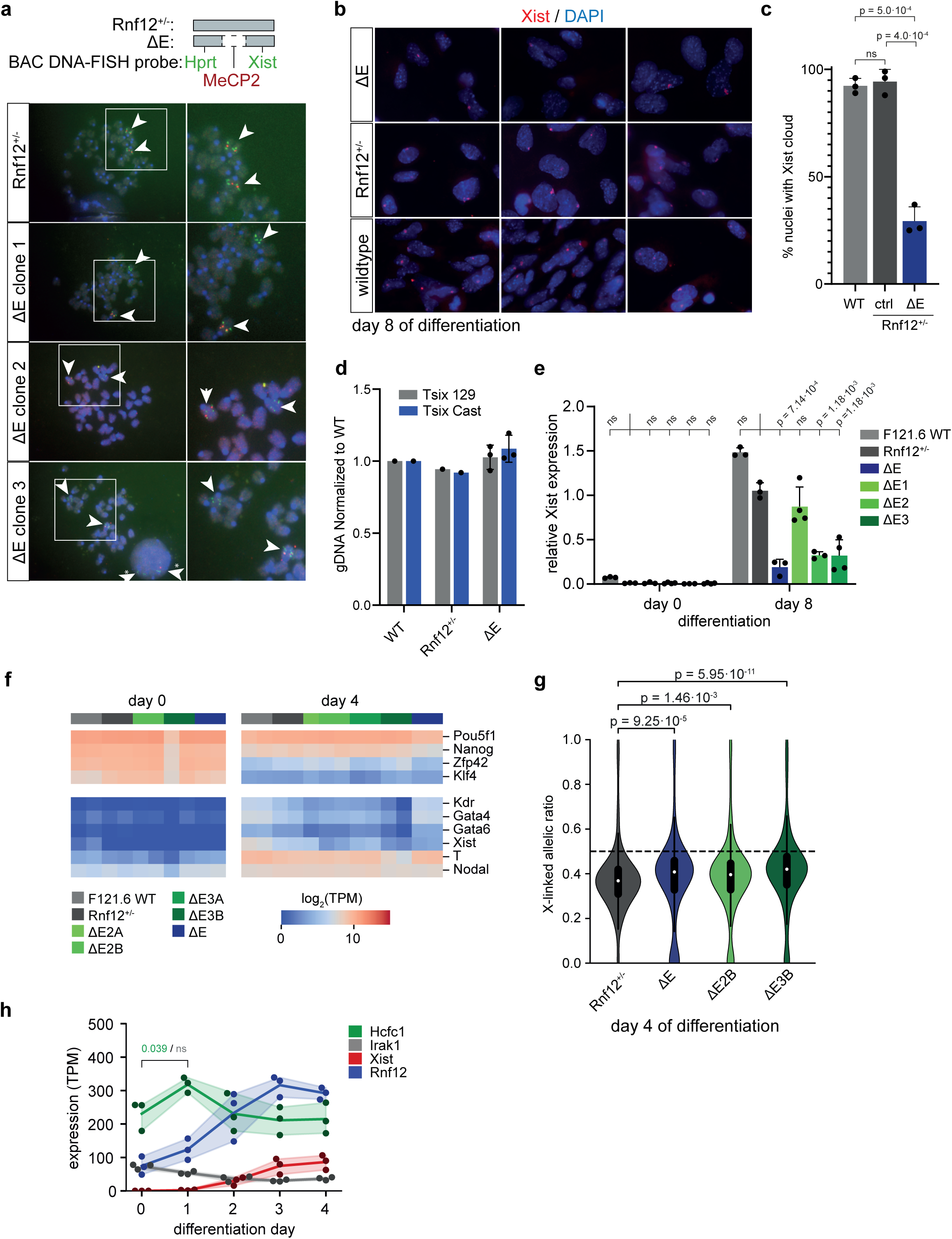
Deletions containing *Hcfc1* show reduced Xist upregulation but can differentiate. **a**, DNA FISH analysis on metaphase spreads of the parental and three ΔE clones detecting *Hprt* and *Xist* in green, and *Mecp2* in red. Notice red is inside the deleted region. White arrowheads indicate chromosome X. Arrowheads with an asterisk in ΔE clone 3 indicate the signal within an interphase nucleus. **b**, Representative images of *Xist* RNA FISH (red) in WT, *Rnf12^+/−^* and three ΔE clones at day 8 of differentiation. **c**, Quantification of **b**, average number of cells with a *Xist* cloud ± SD, n=3 biological replicas, n=201-231, statistical significance was calculated using a two-sided Welch T-test and corrected for multiple testing using the Benjamini-Hochberg procedure. P-values are indicated, ns = not significant. **d**, Allelic gDNA PCR of *Tsix* in WT, *Rnf12^+/−^* and 3 independent ΔE clones on day 8 of differentiation to confirm the presence of two X chromosomes. Average ± SD, n=1-3 biological replicates. **e**, Relative *Xist* expression in WT, *Rnf12^+/−^*, ΔE, ΔE1, ΔE2 and ΔE3 ESCs at day 0 and 8 of differentiation. A one-sided Welch’s t-test was used to test significant differences between *Rnf12^+/−^* and WT or deletion cell lines. P-values were corrected for multiple testing with the Benjamini-Hochberg method. **f**, Heatmap of gene expression (in TPMs) for pluripotency markers (*Pou5f1*, *Nanog*, *Zfp42*/*Rex1*, *Klf4*) and differentiation markers (*Kdr*/*Flk-1*, *Gata4*, *Gata6*, *T* and *Nodal*) across WT, *Rnf12^+/−^* and ΔE, ΔE2A, ΔE2B, ΔE3A and ΔE3B clones. **g**, Violin plot of allelic ratios of X-linked gene expression in *Rnf12^+/−^*, ΔE, ΔE2B and ΔE3B. The box plots display the median (black line), the interquartile range (box limits) and 1.5x of the interquartile range (whiskers). Dashed line indicates ratio of 0.5 as expected ratio without silencing. P-values were calculated using a two-sided Mann-Whitney U test and corrected for multiple testing using the Benjamini-Hochberg approach. **h**, Expression (TPM) of *Hcfc1* (green), *Irak1* (grey), *Rnf12* (blue) and *Xist* (red) in ΔXic_B6_ ESCs from Pacini et al., 2021^19^. P-values were calculated between day 0 and 1 for Hcfc1 (green) and Irak1 (grey) using one-sided independent t-tests and corrected for multiple testing with the Benjamini-Hochberg method.

**Extended Data Figure 3:**
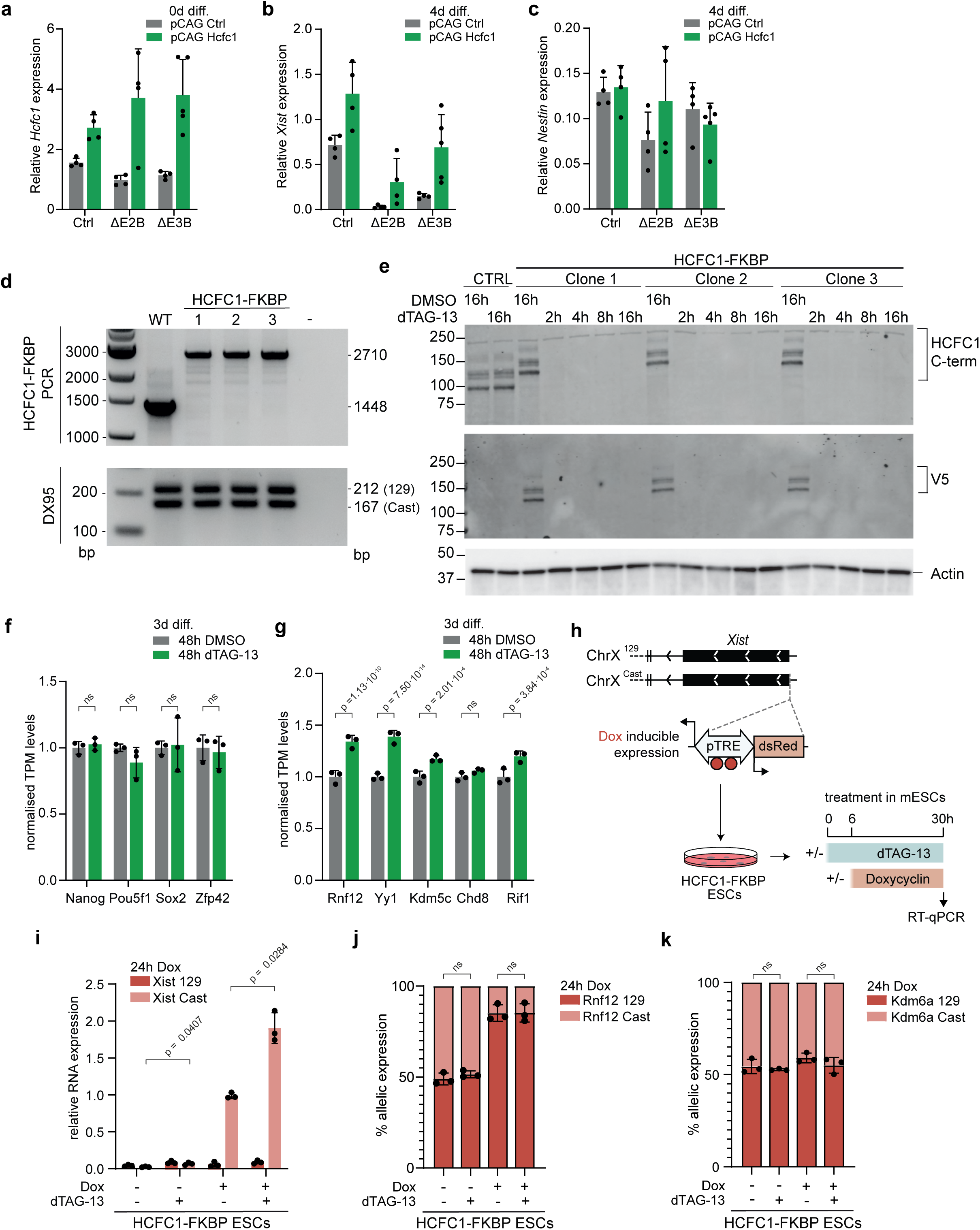
HCFC1 rescue and conditional knockout confirmation. **a**, Relative *Hcfc1* expression in control (*Rnf12^+/−^*), ΔE2B and ΔE3B ESCs upon stable integration, *Hcfc1* expression vs control plasmid in different individual undifferentiated clones; Average ± SD, n=4-5 independent clones. **b**, Relative *Xist* expression in Control (*Rnf12^+/−^*), ΔE2B and ΔE3B upon stable integration, *Hcfc1* expression vs control plasmid at day 4 of differentiation in different individual clones; Average ± SD, n=4-5 independent clones. **c**, Same as in **b**, but for differentiation marker *Nestin*. **d**, PCR analysis of HCFC1-FKBP ESCs, top. DX95 PCR to control for the presence of both X chromosomes, bottom. **e**, HCFC1 C-term and V5 WB in HCFC1-FKBP ESCs after treatment with dTAG-13 or DMSO control in the ESC state. Actin is used as loading control. **f**, Expression of several pluripotency markers at day 3 of differentiation after 48h of dTAG-13 or DMSO treatment. TPM values are normalized to the average expression in the DMSO condition for each gene. P-values were calculated using DESeq2, which corrects for multiple testing using the Benjamini-Hochberg method **g**, Same as in **f**, but for genes involved in *Xist* regulation. **h**, Schematic of HCFC1-FKBP ESCs with a doxycycline-inducible *Xist* promoter and strategy to induce *Xist* expression in the ESC stage in the presence of dTAG-13. **i**, RT-qPCR analysis of *Xist* expression in HCFC1-FKBP ESCs upon 24h of dox treatment and/or 30h dTAG-13 treatment. Average ± SD, n=3 biological replicates. **j**, Allelic expression of X-linked gene *Rnf12* in HCFC1-FKBP ESCs upon 24h Dox and/or 30h dTAG-13 treatment. Average ± SD, n=3 biological replicates. **k**, Allelic expression of XCI escapee *Kdm6a* upon forced *Xist* expression (+Dox) for 24h in the presence of dTAG-13 or DMSO. For (i–k), p-values from two-sided Welch’s t-test between DMSO and dTAG-13 treatment, corrected using the Benjamini–Hochberg procedure.

**Extended Data Figure 4:**
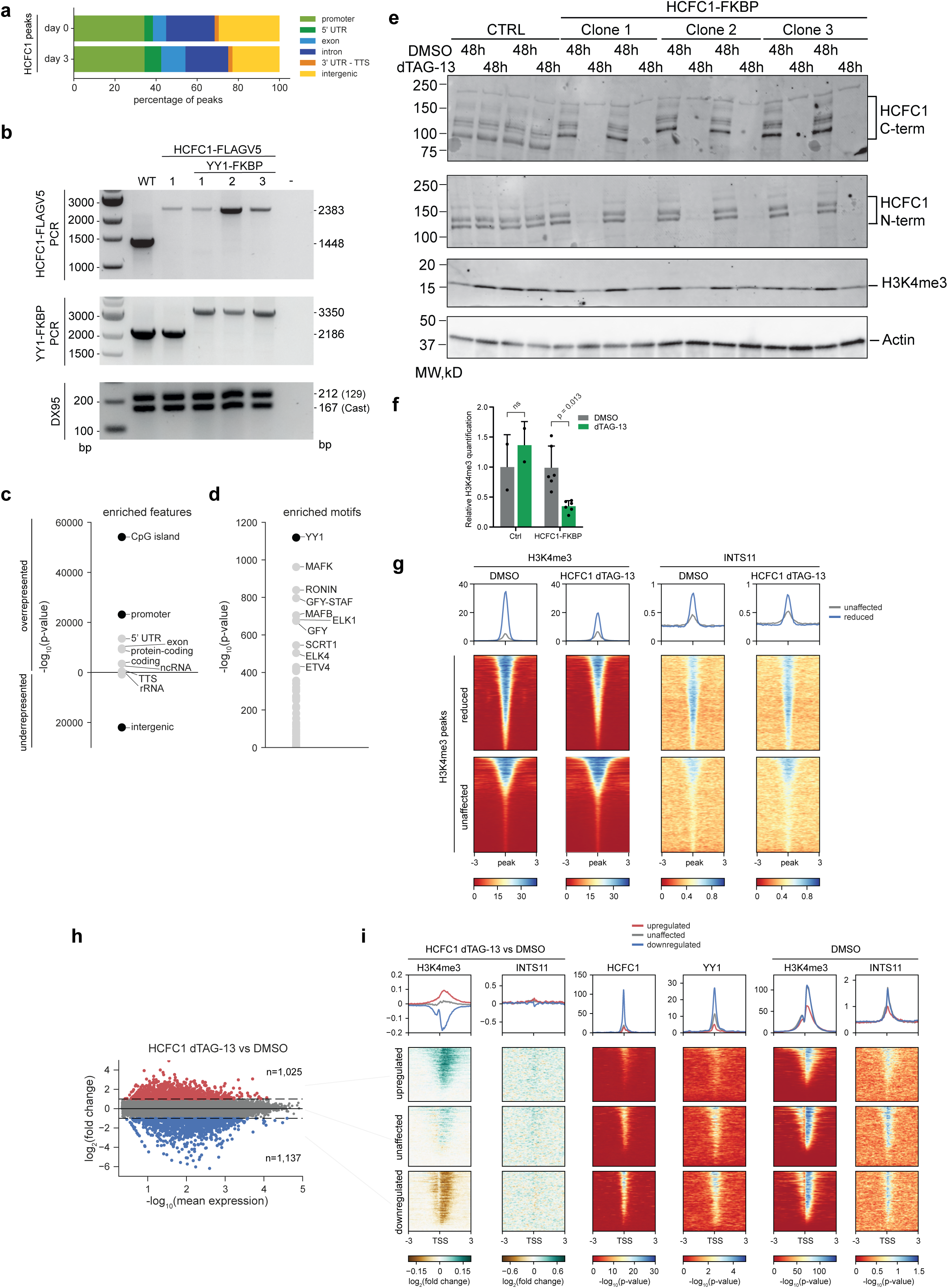
HCFC1 binding sites analysis and genome-wide H3K4me3 and INTS11 recruitment upon dTAG-13-mediated depletion of HCFC1. **a**, Genomic annotation of HCFC1 peaks at day 0 and day 3 of ESC differentiation. **b**, PCR analysis confirming proper integration of HCFC1-V5- and the YY-FKBP-tags and presence of both X chromosomes (DX95). **c**, Feature enrichment analysis at HCFC1 peaks at day 3 of differentiation. Features are grouped by over- and underrepresentation, and the top 10 with the lowest p-values are labeled. **d**, Motif enrichment analysis at HCFC1 peaks at day 3 of differentiation. The 10 most significantly enriched motifs are labeled. **e**, Western blot analysis of H3K4me3 in HCFC1-FKBP cells treated with DMSO or dTAG-13 (48h) at day 3 of differentiation. **f**, Quantification of global H3K4me3 WB signal in Ctrl and HCFC1-FKBP cells treated with DMSO or dTAG-13 (48h) at day 3 of differentiation. ns, not significant, p-values were calculated using a two-sided Welch’s t-test and corrected *via* Benjamini-Hochberg. **g**, Heatmap of H3K4me3 and INTS11 enrichment at H3K4me3 peaks lost upon HCFC1 depletion, and at unaffected peaks (random subset of 4,991 peaks). **h**, MA plot showing gene expression changes upon dTAG-13-mediated depletion of HCFC1. Genes are colored based on significant upregulation (red), downregulation (blue) or no change (grey). **j**, Heatmap of H3K4me3 and INTS11 changes upon HCFC1 depletion, as well as HCFC1 and YY1 binding and H3K4me3 and INTS11 enrichment in the DMSO condition, at promoters of upregulated, unaffected and downregulated genes.

**Extended Data Figure 5:**
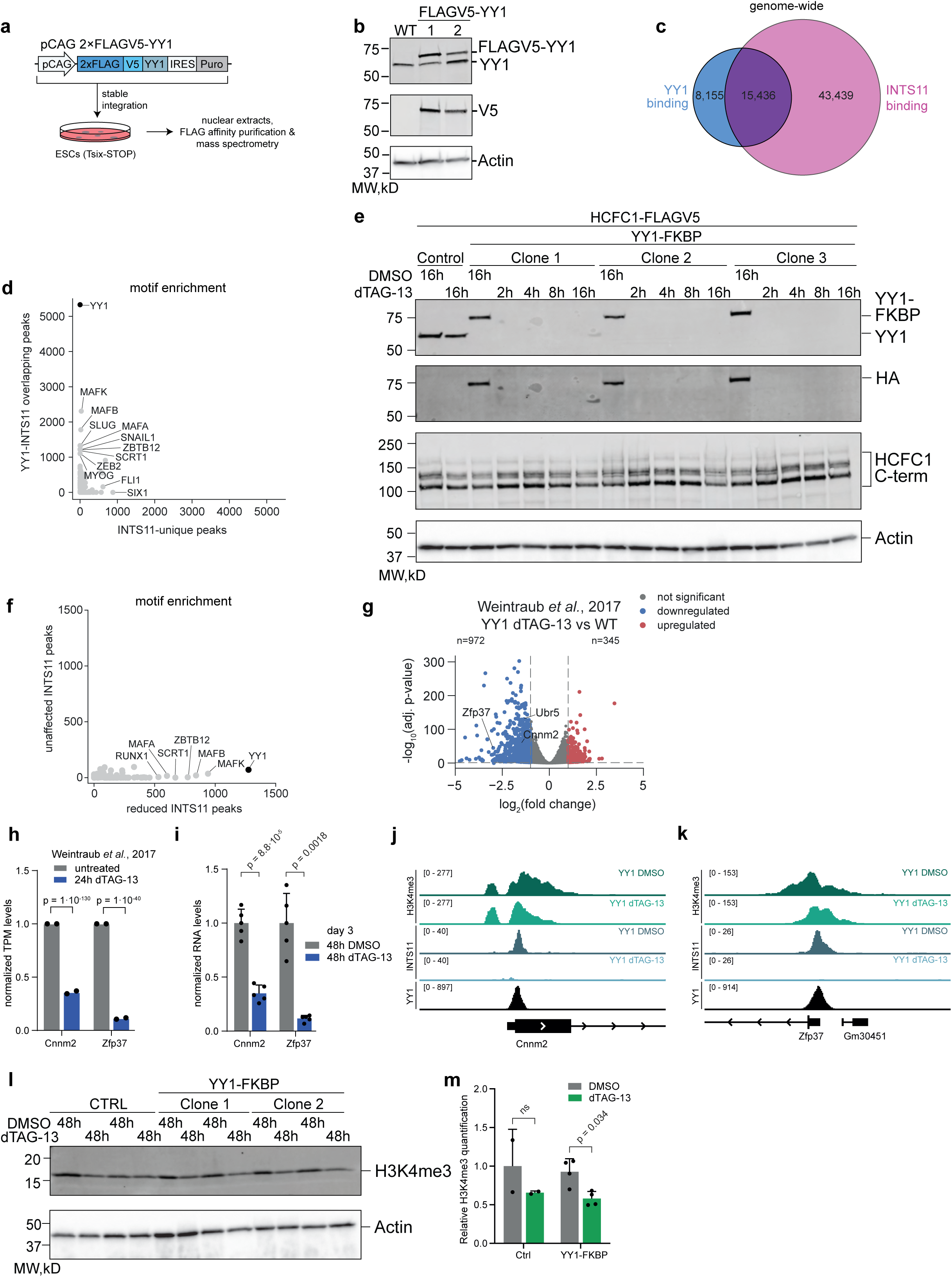
YY1 and INTS11 binding sites analysis. **a**, Generation strategy of the 2xFLAG-V5-Yy1 ESC lines by random integration in XX *Tsix*-STOP ESCs. **b**, Western blotting analysis of YY1 expression in 2xFLAG-V5-YY1 ESCs using an endogenous YY1 or V5 antibody, and Actin as loading control. **c**, Venn diagram showing the genome-wide overlap of YY1 and INTS11 peaks at day 0 of differentiation. **d**, Scatter plot comparing motif enrichment between YY1-INTS11 overlapping peaks and INTS11-unique peaks identified in **c**. Each point represents a motif, plotted by its −log10(p-value) in both analyses, and motifs with the lowest p-values in either condition are labeled. **e**, WB analysis of YY1 degradation upon dTAG-13 treatment vs DMSO for different hours of YY1-FKBP ESCs at day 0 of differentiation. **f**, Scatter plot comparing motif enrichment between reduced and unaffected INTS11 peaks in YY1-FKBP ESCs in the dTAG-13 condition at day 3 of differentiation. Each point represents a motif, plotted by its −log10(p-value) in both analyses, and motifs with the lowest p-values in either condition are labeled. **g**, Volcano plot showing gene expression changes upon YY1 depletion. Genes are colored based on significant upregulation (red), downregulation (blue) or no change (grey). **h**, Expression of *Cnnm2* and *Zfp37* upon 24h YY1 removal in undifferentiated ESCs^33^, average of n=2 biological replicates. TPM values are normalized to the average expression in the DMSO condition for each gene. P-values were calculated using Wald tests and corrected for multiple testing using the Benjamini-Hochberg method, as implemented by DESeq2. **i**, *Cnnm2* and *Zfp37* expression analysis by qRT-PCR in KFBP-YY1 ESCs upon 48h DMSO or dTAG-13 treatment at day 3 of differentiation. A two-sided Welch’s t-test was used to assess significant differences between DMSO- and dTAG-13 treated cells. P-values were corrected for multiple testing using the Benjamini-Hochberg method. **j**, Views of ChIP-seq tracks for H3K4me3 (green) and INTS11 (blue) in YY1-depleted ESCs versus its DMSO control at day 3 of differentiation, and YY1 ChIP-seq track in female ESCs at day 0 (black), around the *Cnnm2* promoter region. **k**, Same as **j**, for *Zfp37.* **l**, WB analysis of YY1 and H3K4me3 of YY1-FKBP ESCs differentiated for 3 days and treated with dTAG-13 or DMSO for the last 48h. **m**, Quantification of global H3K4me3 WB signal in Ctrl and YY1-FKBP cells treated with DMSO or dTAG-13 (48h) at day 3 of differentiation. ns, not significant. P-values were calculated using a two-sided Welch’s t-test and corrected *via* Benjamini-Hochberg.

